# SMRT sequencing yields the chromosome-scale reference genome of tea tree, *Camellia sinensis* var. *sinensis*

**DOI:** 10.1101/2020.01.02.892430

**Authors:** Qun-Jie Zhang, Wei Li, Kui Li, Hong Nan, Cong Shi, Yun Zhang, Zhang-Yan Dai, Yang-Lei Lin, Xiao-Lan Yang, Yan Tong, Dan Zhang, Cui Lu, Chen-feng Wang, Xiao-xin Liu, Wen-Kai Jiang, Xing-Hua Wang, Xing-Cai Zhang, Zhong-Hua Liu, Evan E. Eichler, Li-Zhi Gao

## Abstract

Tea is the oldest and most popular nonalcoholic beverage consumed in the world. It provides abundant secondary metabolites that account for its diverse flavors and health benefits. Here we present the first high-quality chromosome-length reference genome of *C. sinensis* var. *sinensis* using long read single-molecule real time (SMRT) sequencing and Hi-C technologies to anchor the ∼2.85-Gb genome assembly into 15 pseudo-chromosomes with a scaffold N50 length of ∼195.68 Mb. We annotated at least 2.17 Gb (∼74.13%) of repetitive sequences and high-confidence prediction of 40,812 protein-coding genes in the ∼2.92-Gb genome assembly. This accurately assembled genome allows us to comprehensively annotate functionally important gene families such as those involved in the biosynthesis of catechins, theanine and caffeine. The contiguous genome assembly provides the first view of the repetitive landscape allowing us to accurately characterize retrotransposon diversity. The large tea tree genome is dominated by a handful of Ty3-*gypsy* long terminal repeat (LTR) retrotransposon families that recently expanded to high copy numbers. We uncover the latest bursts of numerous non-autonomous LTR retrotransposons that may interfere with the propagation of autonomous retroelements. This reference genome sequence will largely facilitate the improvement of agronomically important traits relevant to the tea quality and production.

## Introduction

Tea is the oldest (since 3000 BC) and most popular nonalcoholic beverage in the world. It is one of the most economically important crops grown in China, India, Sri Lanka, and Kenya with approximately 3.0 million metric tons of dried tea produced annually (Chen et al., 2007; Soni et al., 2015). Besides a wealth of health benefits, it has also long affected the culture, health, medicine, and trade around Asia, and even the world (Banerjee, 1992; Liu et al., 2019; Mondal et al., 2004). The tea tree *Camellia sinensis* L. O. Kuntze, a member of the genus *Camellia* in the Theaceae family, is the source of commercially grown tea for nearly 5,000 years, (Taniguchi et al., 2014; Wheeler and Wheeler, 2004). Besides other wild species of the section *Thea* cultivated in small quantities, such as *C. taliensis, C. grandibracteata, C. sinensis* var. *dehungensis, C. sinensis* var. *pubilimba* and *C. ptilophylla*, the most widely grown tea tree (*C. sinensis*) includes the two major varieties: *C. sinensis* var. *sinensis* (*CSS*; Chinese type) and *C. sinensis* var. *assamica* (*CSA*; Assam type) (Ming and Bartholomew, 2007). *CSS* is a slow-growing shrub with small leaves and can tolerate cold climates, making it adaptable to a broad geographic range, and has become the most popular elite tea tree cultivar in China (∼67%) (Willson and Clifford, 2012). *CSA* is quick-growing with large leaves and mainly cultivated in tropical and subtropical regions, due to high sensitivity to cold weather, such as Yunnan Province in China and India (Willson and Clifford, 2012).

The health-promoting functions of tea are attributable to the presence of bioactive compounds with strong antioxidant properties (Liu et al., 2019). Among a large number of metabolites, the most characteristic are catechins (a subgroup of flavan-3-ols), theanine (γ-glutamylethylamide) and caffeine. Catechins mainly confer an astringent taste to tea, theanine contributes to the umami and sweet tastes of tea infusions, while caffeine offers a bitter taste (Narukawa et al., 2008). The ratio of phenol to ammonia usually forms a basis for the choice of tea processing procedures. *CSA* is usually processed into black tea for its high content of catechins, and catechins are polymerized to theaflavins and thearubigins by a “fermentation” that leads to oxidation of the catechins, while *CSS* can be processed into green tea, which retain the astringency and the antioxidant activity of catechins (Li et al., 2013).

Considering the tremendously economic importance of the tea tree, there have been constant efforts to explore the genetic basis of the biosynthesis of natural metabolites that determine health benefits as well as the formation of diverse tea flavors (Li et al., 2015; Liu et al., 2019; Shi et al., 2011; Xia et al., 2017). Modern improvements to biotic resistance and abiotic tolerance in the tea tree-breeding programs are necessary not only for tea quality and yields but also for the consumer safety on tea from harmful organisms and pesticide residues. The progress in tea tree genomics is an essential solution, which largely relies on the completion of a high-quality reference genome sequence. We released the first draft genome sequence of *C. sinensis* var. *assamica* cv. *Yunkang-10* (*CSA-YK10*) using whole-genome shotgun Illumina sequencing technology, providing the first insights into the genomic basis of tea flavors and global adaptation (Xia et al., 2017). The second tea tree draft genome was followed by sequencing *C. sinensis* var. *sinensis* cv. *Shuchazao* (*CSS-SCZ*) using the same sequencing platform and then filling gaps with PacBio long reads (Wei et al., 2018). However, obtaining a high-quality tea tree genome assembly remains a great challenge, because short Illumina reads and even hybrid assembly strategies have always been problematic to *de novo* assemble any complex plant genome having highly heterozygous and repetitive DNA sequences. As one of the longest transposable elements (TEs), long terminal repeat (LTR) retrotransposons are insoluble for short Illumina reads. However, the abundance makes them serve as an important driver of the genome size variation in flowering plants (Piegu et al., 2006; Vitte and Panaud, 2005). LTR retrotransposons in the tea tree genome, for example, represent the majority (∼67.21%) of the *CSA-YK10* genome assembly (Xia et al., 2017).

Here, we present a highly contiguous tea tree genome assembly of an elite tea tree cultivar, *C. sinensis* var. *sinensis* cv. *Biyun* (*CSS-BY*), based on long-read single-molecule real-time sequencing (SMRT) and Hi-C technologies. We obtain accurate genomic information for almost all gene families, such as those involved in the biosynthesis of flavonoids, theanine and caffeine that contribute to tea flavors and health benefits, providing novel insights into the evolution of non-autonomous long terminal repeat (LTR) retrotransposons that affect the increasing of the large e genome size.

## Results and Discussion

*De novo* sequencing and assembling the highly heterozygous tea tree genome have long been challenging as a result of its self-incompatible nature (Xia et al., 2017). We employed the Illumina short-read technology with paired-end libraries on the HiSeq X Ten sequencing platform to screen 12 representative tea tree cultivars. This generated raw sequence data sets of 1,679.6 Gb, thus yielding approximately 508.76-fold high-quality sequence coverage for all varieties (**Supplementary Table 1**). We thus selected the commercial variety (*CSS-BY*) for long-read genome sequencing due to its relatively low heterozygosity (1.22%). We estimated that the genome size of *CSS-BY* is 3.25 Gb using 17-mer analysis (**Supplementary Figure 1**; **Supplementary Table 1**). We performed a whole-genome shotgun sequencing (WGS) analysis with the PacBio SMRT sequencing platform. This generated clean sequence data sets of 417.95 Gb with an average read length of 11.9 Kb and yielded approximately 127.66-fold coverage (**Supplementary Table 2**). Then, ∼282.94 Gb high-quality next-generation sequencing (NGS) data with 86.42-fold genome coverage using the Illumina HiSeq X Ten platform were employed to polish the assembled genome (**Supplementary Table 1**). A total of 909,454,810 Hi-C reads (**Supplementary Table 3**) were used to connect pseudo-chromosomes by using LACHESIS (Burton et al., 2013) and JUICEBOX (Durand et al., 2016; Robinson et al., 2018). This final assembly of the tea tree genome was ∼2.92 Gb, accounting for ∼89.85% of the estimated genome size; ∼2.85-Gb of the genome assembly (∼97.88%) was anchored into 15 pseudo-chromosomes (**Figure 1; Table 1; Supplementary Figure 2; Supplementary Tables 4-6**). The assembly comprised 13,006 contigs with a contig N50 length of 625.11 Kb, ∼9.32 times longer than the previously reported genome assembly of *C. sinensis* var. *sinensis* cv. *Shuchazao* (*CSS-SCZ*) (∼67.07 Kb) that was assembled by Illumina reads and then filled gaps with PacBio single-molecule long reads (Wei et al., 2018) and 31.32 times longer than *C. sinensis* var. *assamica* cv. *Yunkang-10* (*CSA-YK10*) (19.96 Kb) that was assembled by Illumina reads only (Xia et al., 2017) **(Table 1)**. The assembly was comprised of 4,153 scaffolds with a scaffold N50 length of 195.68 Mb, ∼140.78 times longer than the previously reported genome assembly of *C. sinensis* var. *sinensis* cv. *Shuchazao* (*CSS-SCZ*) (∼1.39 Mb), which was assembled by Illumina reads and then gaps filled with PacBio SMRT long reads (Wei et al., 2018) **(Table 1)**. The lengths of 15 chromosomes of the *CSS-BY* genome ranged from ∼253 Mbp (Chr01) to ∼128 Mbp (Chr15) with an average size of ∼190 Mbp (**Figure 1**; **Supplementary Table 6**). Our results showed that 98.22% of NGS reads could be unambiguously represented with an expected insert size distribution spanning 98.44% of the assembled genome, indicating a high confidence of genome scaffolding (**Supplementary Table 7**). We further applied CEGMA (Core Eukaryotic Gene Mapping Approach) (Parra et al., 2007) to assess the quality of the genome assembly. CEGMA assessment showed that 227 of 248 core eukaryotic genes (91.53%) were completely assembled, and only 9 genes (3.63%) were partially presented (**Supplementary Table 8**). We finally assessed core gene statistics using BUSCO (Benchmarking Universal Single-Copy Orthologs) (Simão et al., 2015) to verify the sensitivity of gene prediction, completeness and propriety of removing redundant sequences of the genome assembly. Our predicted genes resolved 88.13% of complete BUSCOs and only 3.68% of fragmented BUSCOs in the Embryophyta lineage (**Supplementary Table 9**).

**Table 1.** Global statistics for the assembly and annotation of the two *Camellia sinensis* var. *sinensis* genome assemblies.

|  | *CSS-BY* | *CSS-SCZ* |
| --- | --- | --- |
| **Assembly** |  |  |
| Estimated genome size (Gb) | 3.25 | 2.98 |
| Total length of scaffolds (Gb) | 2.92 | 3.14 |
| Coverage of the assembled sequences (%) | 89.85 | 1.05 |
| Scaffold number | 4,153 | 14,051 |
| N50 of scaffolds (Mb) | 195.68 | 1.39 |
| N50 of contigs (Kb) | 625.11 | 67.07 |
| GC content of the genome (%) | 38.24 | 37.84 |
| **Annotation** |  |  |
| No. of predicted protein-coding genes | 40,812 | 33,932 |
| Average gene length (bp) | 6,263 | 4,053 |
| Average exon length (bp) | 251 | 259 |
| Average exon per gene | 5.2 | 3.3 |
| Mean intron length (bp) | 1,168 | 1,408 |
| tRNAs | 659 | 597 |
| rRNAs | 2,845 | 2,838 |
| snoRNAs | 471 | NA |
| snRNAs | 207 | 416 |
| miRNAs | 139 | 355 |
| Masked repeat sequence length (Mb) | 2,165 | 2,008 |
| Percentage of repeat sequences (%) | 74.13 | 64.78 |

**Figure 1.**
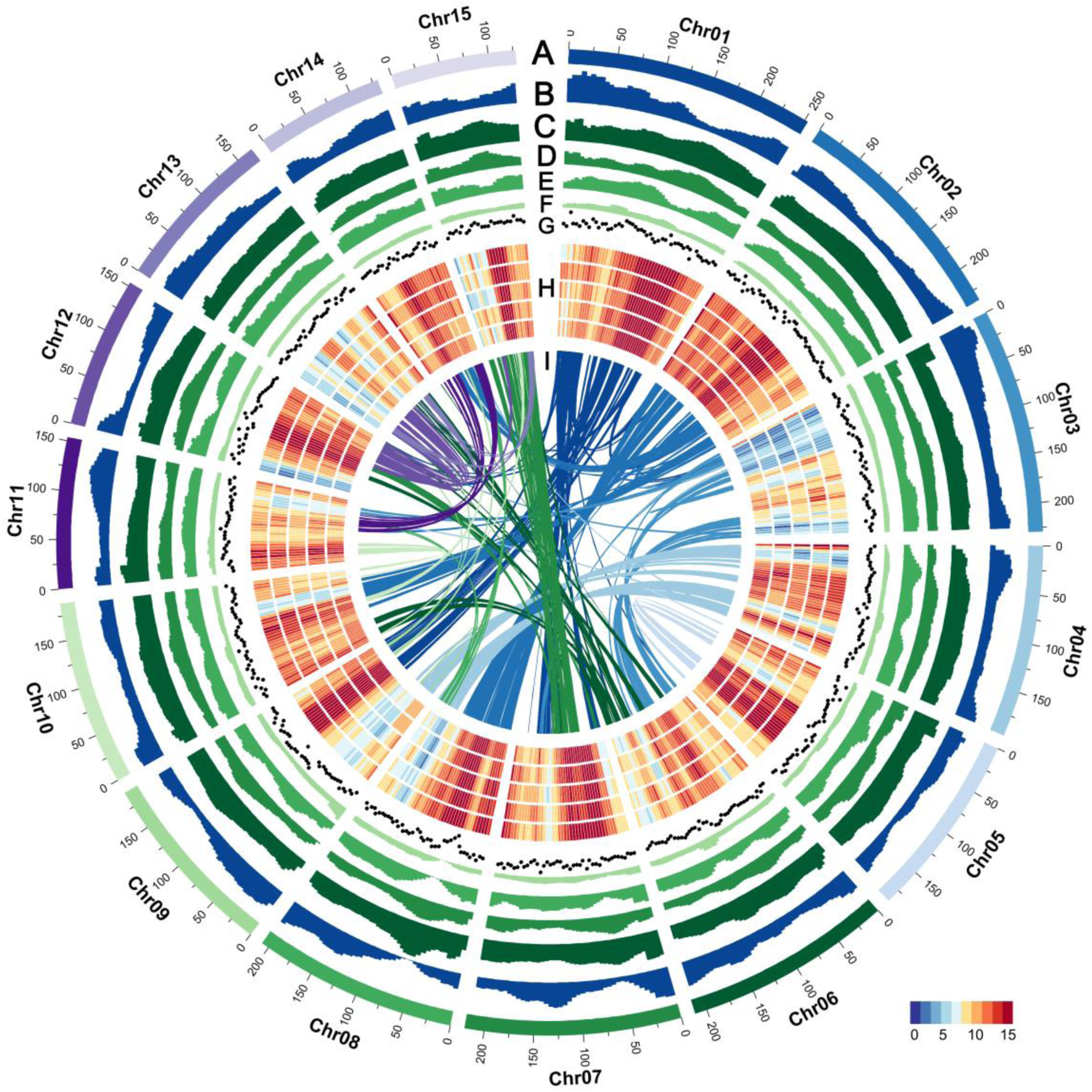
The genome features of *C. sinensis* var. *sinensis* cv. *Biyun*. (**A**) Circular representation of the 15 pseudochromosomes. (**B**) The density of genes. (**C**) The distribution of TEs. (**D**) The distribution of Ty3-*gypsy* LTR-RTs. (**E**) The distribution of Ty1-*copia* LTR-RTs. (**F**) The distribution of DNA TEs. (**G**) The density of SSRs. (**H**) The density of transcript expression for young leaf (YL), tender shoot (TS), flower bud (FB), fruit (FR) and stem (ST) from outside to inside. (**I**) Genomic synteny.

**Figure 2.**
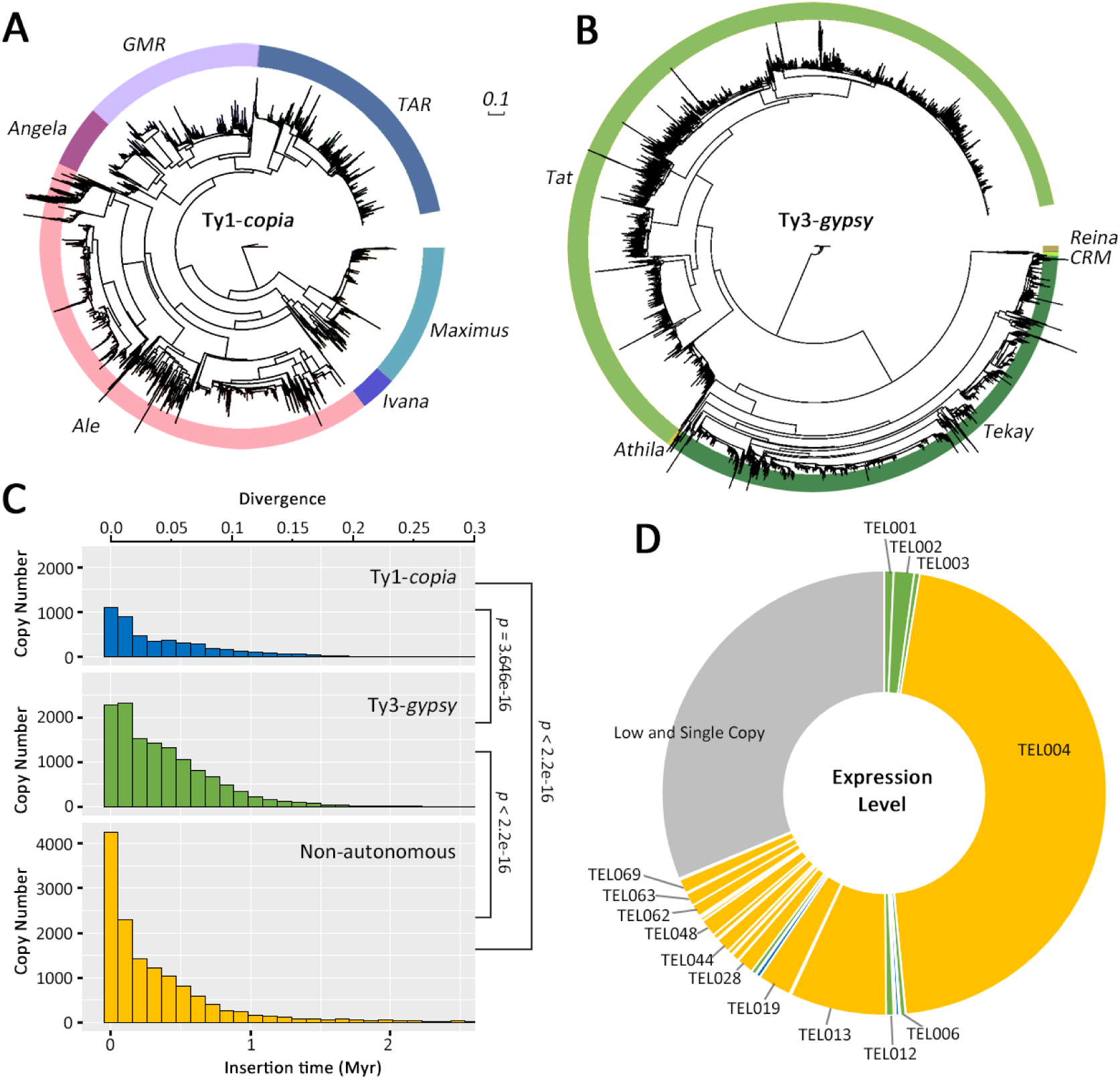
The evolutionary landscape of LTR retrotransposons in the *C. sinensis* var. *sinensis* cv. *Biyun* genome. The neighbor-joining and unrooted phylogenetic trees were constructed based on 4,630 Ty1-*copia* (**A**) and 13,172 Ty3-*gypsy* (**B**) aligned sequences corresponding to the RT domains without premature termination codon. (**C)** Insertion times of Ty1-*copia* (blue), Ty3-*gypsy* (green) and non-autonomous (yellow) LTR retrotransposons; The insertion times for LTR retrotransposons were calculated by the formula of T = K/2*r*. T: insertion time; r: synonymous mutations/site/Myr; K: the divergence between the two LTRs. A substitution rate of 5.62×10 ^-9^ per site per year (Huang et al., 2013; Shi et al., 2010) was used to calculate the insertion times. (**D**) Expression levels calculated by transcripts read count of LTR retrotransposon families. All transcripts from five tissues were by using TopHat and Cufflinks to classify the LTR retrotransposons related transcripts into different LTR families by BLAST. Then, reads number of each LTR retrotransposon family were counted by HTSeq.

We annotated approximately 2,164.89 Mb (∼74.13%) of repetitive sequences in the *CSS-BY* genome assembly (**Supplementary Figure 3A; Supplementary Table 10**). The total content of repetitive sequences in the *CSS-BY* genome is apparently larger than the formerly reported *CSS-SCZ* genome assembly (∼64.77%, 2,008.28 Mb) (Wei et al., 2018), consistent with a more comprehensive *de novo* assembly of genomic regions containing highly repetitive sequences using long PacBio reads (**Supplementary Figure 3A**; **Supplementary Table 10**). We annotated 32,367 full-length LTR retrotransposons in the *CSS-BY* genome, which are ∼2.5 times more abundant than *CSS-SCZ* (13,119) **(Supplementary Figure 3D**; **Supplementary Table 11)**. All these results together demonstrate that, besides the possibility of genome size variation among tea tree cultivars, high-quality PacBio-only *CSS-BY* genome assembly, has improved the detection of repeat content when compared to the previous NGS-based genome assemblies (*CSA-YK10* and *CSS-SCZ*) (**Supplementary Table 10**).

**Figure 3.**
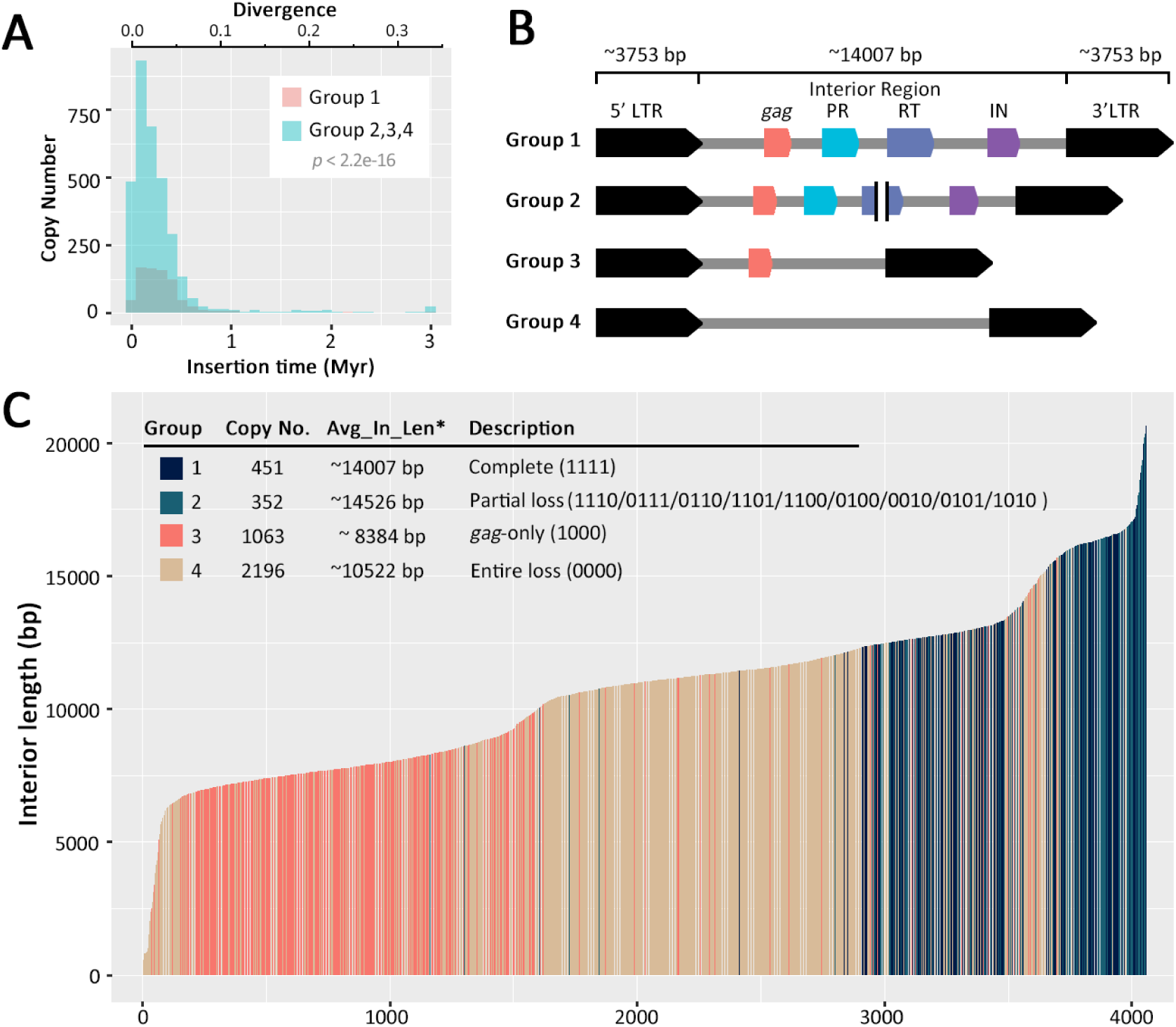
Evolutionary dynamics of the top retrotransposon family in the *C. sinensis* var. *sinensis* cv. *Biyun* genome. (**A**) Insertion times of LTR retrotransposons.(**B**) Structural features of the four groups of the top TEL001 retrotransposon family. (**C**) Length distribution of interior regions of the TEL001 retrotransposon family. Group 1 (dark blue) stands for complete retroelements containing *gag* and *pol* (PR, RT and IN) genes; Group 2 (steal blue) stands for any one or more partial open reading frames (ORFs) (but not all) that are not fully encoded; Group 3 (orange) for retroelements with only *gag*; and Group 4 (camel) for retroelements without any ORFs. The four digits in the ‘Description’ represent the *gag* and *pol* (PR, RT and IN) genes in LTR retrotransposon, respectively. The number ‘1’ represents the presence of a Pfam annotation, and ‘0’ indicates the absence. For example, ‘1110’ means that the LTR retrotransposon contain a *gag*, PR and RT without IN.

We combined *ab initio* prediction and transcriptome sequence alignments from RNA-seq data for five tissues, including young leaf (YL), tender shoot (TS), flower bud (FB), fruit (FR), and stem (ST) to annotate protein-coding genes (**Supplementary Tables 12-13**). Using rigorous filter parameters, we totally predicted 40,812 protein-coding genes **(Table 1**), of which 34,722 (85.08%) genes were supported by transcriptome-based evidence (**Supplementary Table 14**). The average gene length and exon number were 6,263 bp and 5.2 per gene, which are much higher than those in *CSS-SCZ* with 4,053 bp and 3.3 per gene, respectively (**Table 1; Supplementary Figure 4**). Of them, 95.64%, 78.39%, 73.17%, 17.98%, 60.12% and 21.62% could be functioned with InterProScan (Jones et al., 2014), SwissProt (Boeckmann et al., 2003), Pfam (Finn et al., 2013), KEGG (Kanehisa and Goto, 2000), GO (Ashburner et al., 2000) and TmHMM (Möller et al., 2001) databases, respectively (**Supplementary Table 14**; **Supplementary Figures 5-6**).

The annotation of noncoding RNA (ncRNA) genes yielded 659 transfer RNA (tRNA), 2,845 ribosomal RNA (rRNA), 471 small nucleolar RNA (snoRNA), 207 small nuclear RNA (snRNA) and 139 microRNA (miRNA) genes (**Supplementary Table 15**). For miRNAs, a total of 2,016 miRNA target sites were predicted using psRNATarget server (**Supplementary URLs**). The annotation using GO and KEGG databases showed that these miRNA target genes were enriched in signaling (GO:0023052), catalytic activity (GO: 0003824) and binding (0005488) (**Supplementary Figure 7**), and were enriched in genetic information processing, organismal systems,carbohydrate metabolism and environmental information processing (**Supplementary Figure 8**). We annotated 1,673,577 simple sequence repeats (SSRs), which may provide valuable genetic markers to assist future tea tree genetic improvement programs (**Supplementary Table 16**; **Supplementary Figure 9**).

Comparative analyses of the *CSS-BY* and *CSS-SCZ* genome assemblies surprisingly detected only 16,313 collinear genes (21.80% in a total of 74,822 genes) at the scaffold level using MCScanX (Wang et al., 2012) (**Supplementary Table 17)**. Such an unbelievably low genome collinearity between the two varieties of *C. sinensis* var. *sinensis* (*CSS-BY* and *CSS-SCZ*) hints that at least one of the two genome assemblies are most likely to be incorrectly assembled. Statistics of assembled contigs revealed remarkably higher genome assembly contiguity of the *CSS-BY* assembly than the *CSS-SCZ* assembly, evidenced by much fewer numbers of contigs with longer sizes (**Table 1; Supplementary Figure 10**). Of note, the *CSS-BY* assembly has the longest contig at ∼3.91 Mb, while the longest contig for *CSS-SCZ* is only ∼0.54 Mb (**Supplementary Figure 11**). The top 136 longer contigs, accounting for ∼10% (∼300 Mb) of the *CSS-BY* assembly, corresponded to 1,355 contigs in the *CSS-SCZ* assembly (**Supplementary Figure 11**). Assembly quality was further evaluated by comparative genomic analyses of the selected homologous regions between the *CSS-SCZ* and *CSS-BY* assemblies (**Supplementary Figure 3B**; **Supplementary Figure 12**). Using MUMmer 4 (Delcher et al., 2003), we annotated an exemplar contig, ctg7832 (Chr01:143191430..146616428) (3,424,999 bp) from our *CSS-BY* genome assembly, which corresponded to 35 contigs with an average length of ∼64,705 bp derived from up to 14 scaffolds (21,664,538 bp) in the *CSS-SCZ* assembly (**Supplementary Figure 3B**). The annotation of this *CSS-BY* contig yielded 21 genes, which linked to 19 genes from 15 contigs of the *CSS-SCZ* assembly (**Supplementary Figure 3B**). We also found ctg7832 to be exceedingly abundant in long, high-quality Ty3-*gypsy*-like retrotransposons (2,159,435 bp, 63.05%) with a high quality, particularly containing rather young retroelements (e.g., *Tekay*, 20.62%) in the *CSS-BY* assembly (**Supplementary Figure 3C**). Our results suggest that, compared to the fragmented draft *CSS-SCZ* assembly with limitations of low contiguity of contigs and poor assembly scaffolding, the SMRT sequencing and assembly strategy has produced a *CSS-BY* assembly of superior contiguity containing accurate long-range information, such as recently generated long repeat sequences.

A major motivation for *de novo* tea tree genome assembly is the identification of accurate information of functionally important gene families involved in the biosynthesis of secondary metabolites, such as catechins, theanine and caffeine. With this high-quality genome assembly of *C. sinensis* var. *sinensis* on hand, as a case study, we annotated all 23 gene families encoding enzymes potentially involved in catalyzing reactions of flavonoid, theanine, and caffeine pathways. Our results showed that, besides the four gene families (*UGT84A, GS/TS, GOGAT* and *AMPDA*) with the same copy number between *CSS-BY* and *CSS-SCZ*, up to fifteen gene families (*PAL, C4H, 4CL, CHI, F3H, F3’H, F3’5’H, DFR, FLS, LCR, ANS, ADC, GDH, IMPDH* and *NMT*) had more members in *CSS-BY* than *CSS-SCZ*. Phylogenetic analyses of the annotated genes among three tea tree and kiwifruit genome assemblies strongly support the reality, confirmed by high levels of gene expression for most of novel genes (FPKM ≥ 1) (**Supplementary Tables 19-21**; **Supplementary Figures 13-16**). However, fewer copy numbers were annotated in *CSS-BY* than *CSS-SCZ* for four gene families (*CHS, ANR, SCPL1A* and *SAMS*) (**Supplementary Table 18**). Taking *SCPL1A*, for example, we only annotated 10 members in *CSS-BY* but 22 in *CSS-SCZ*. A phylogenetic analysis indicates that some branches in *CSS-SCZ* had more copies than *CSS-BY*, which had low expression levels (FPKM < 1) (Wei et al., 2018) (**Supplementary Figures 13 and 16**; **Supplementary Table 19**). This does not exclude the occurrence of false positives of redundant genes caused by short reads generated from the Illumina sequencing platform for a highly heterozygous tea tree genome. Our results suggest that the long-read *CSS-BY* genome assembly has undoubtedly promised a reliable annotation of almost all gene families in tea tree.

The long reads generated for the SMRT-based *CSS-BY* genome assembly guarantee to identify almost all transposable elements (TEs), revealing the highly repetitive nature of the tea tree genome (**Figure 1; Supplementary Figure 3A**) and providing the opportunity to understand how the abundance of LTR retrotransposons has contributed to its large genome size. Ty3-*gypsy* LTR retrotransposon elements dominate the genome with ∼34.11% (∼996.15 Mb) of the assembled sequence length, ∼7.11-fold larger than Ty1-*copia* LTR retrotransposon families (∼140.11 Mb; ∼4.80%), and ∼2.03-fold larger than non-autonomous LTR retrotransposon families (∼490.84 Mb; ∼16.81%) (**Supplementary Table 10**; **Supplementary Figure 3A**). The long reads generated for the SMRT-based *CSS-BY* genome assembly guarantee to identify almost all full-length transposable elements, making us the first time to obtain the repetitive evolution history of the genome. To track the evolutionary past of LTR retrotransposons we further classified all full-length LTR retrotransposons into 8,844 families, of which the top 111 families with more than 10 copies contained 75% full-length LTR retrotransposons and occupied 36.47% of the genome, 328 comprised 2-9 copies, and 8,405 were single-copy families (**Supplementary Table 22**). A total of 13,172 Ty3-*gypsy* and 4,630 Ty1-*copia* RT sequences were extracted to construct phylogenetic trees (**Figure 2A** and **2B**), yielding 11 lineages, consistent with previous results (Hřibová et al., 2010; Llorens et al., 2009; Vitte et al., 2007; Wicker and Keller, 2007). *Tat* and *Tekay* accounted for the 98% of Ty3-*gypsy* full-length LTR retrotransposons, indicating a massive expansion during tea tree genome evolution (**Figure 2B**; **Supplementary Figure 20**), while *Ale, TAR, GMR, Maximus Angela*, and *Ivana* of Ty1-*copia* all retained full-length LTR retrotransposons suggesting ing that Ty1-*copia* has always experienced a long and slow amplification history (**Figure 2A**; **Supplementary Figure 20**). The repetitive nature of the tea tree genome is determined by a handful of LTR retrotransposon families with extremely high copy numbers, for example, the amplification of *Tat* (∼671.13Mb; ∼22.98%) and *Tekay* (∼303.84 Mb; ∼10.41%) of Ty3-*gypsy* has largely contributed to the large tea tree genome size (**Supplementary Figure 3B**). Of them, incessant bursts of the *Tat* lineage predominantly came from eight (*TEL001, TEL002, TEL003, TEL006, TEL007, TEL008, TEL011* and *TEL012*) of the top 12 families resulted in ∼50% of full-length LTR retrotransposons that accounted for ∼29.63% of this genome assembly (**Supplementary Table 22**). The largest family *TEL001*, for instance, contained 4,062 full-length LTR retrotransposons with the longest average length of 18,204 bp, contributing most to the genome size (∼18.27%) (**Supplementary Figure 3E and 3B; Supplementary Table 22**).

The availability of the best *CSS* genome assembly of *CSS-BY* so far permits us to investigate how LTR retrotransposons evolve in the tea tree genome. We combined RT sequences from the two major varieties of tea tree, *CSS-BY* and *CSA-YK10*, by adding 4,579 Ty3-*gypsy* and 1,406 Ty1-*copia* RT sequences from *CSA-YK10* (**Supplementary Figure 17**). Our results showed that they may have experienced a similar evolutionary history except that considerably large numbers of retrotransposons (e.g., *Tat* and *Tekay* lineages) were detected in the SMRT-based *CSS-BY* genome assembly.

The resulting 32,367 full-length LTR retrotransposons account for nearly 18.5% of the assembled sequence length and allow us to further trace the very recent evolutionary history of LTR retrotransposons and evolutionary dynamics of the tea tree genome size (**Supplementary Table 10**; **Supplementary Table 22**). The failure to assemble the recently generated retrotransposons by using Illumina short reads for the *CSS-SCZ* genome assembly is evidenced by the detection of a small portion of full-length LTR retrotransposons inserted within the past 1 Myr (**Supplementary Figures 18-19**). In comparison, the SMRT-based *CSS-BY* genome assembly enables us for the first time to date the fairly recent insertion events of LTR retrotransposons, which are necessary to present a clear dynamic history of retrotransposon bursts in the genome. The expansion of Ty3-*gypsy* retrotransposon families makes the genome currently predominate (**Supplementary Figure 3DE**), such as *Tat* members(∼22.98% of the assembled genome)of the Ty3-*gypsy* lineage, which have rapidly amplified during the last 1 Myr, and then rapidly declined in recently (**Supplementary Figures 20-21**). Besides the largest family *TEL001* that increased to a total of 4,062 full-length LTR retrotransposons (∼6.2-fold more than those annotated in the Illumina-based *CSS-SCZ* genome assembly) during the last 1 Myr (**Supplementary Figures 3D and 3E**; **Figure 3A; Supplementary Table 22**), a small number of other multi-copy families belonging to the *Tat* lineage (e.g., *TEL002, TEL003, TEL006, TEL007, TEL008, TEL011* and *TEL012*) have also expanded to large quantities that dominate the *CSS-BY* genome. The finding is consistent with the rapid growth of the *TL001* family in *CSA-YK10* (Xia et al., 2017); the top family was further classified into *TEL001, TEL002, TEL003, TEL006, TEL007* and *TEL008* according to sequence divergence of LTRs (**Supplementary Table 22**). We surprisingly observed that the *Tekay* lineage of Ty3-*gypsy* (e.g., *TEL005, TEL021* and *TEL022*) and nonautonomous LTR retrotransposon families (e.g., *TEL004* and *TEL010*) (**Supplementary Figures 20-21**; **Supplementary Table 22**), accounting for ∼10.41% and ∼16.81% of the genome, respectively (**Supplementary Figure 3A**), were in turn predominant to recently affect dynamic variation of the genome size (**Supplementary Figure 3E**). The retrotransposon abundance is expectedly governed by recent activities of multi-copy LTR retrotransposon families, but it is of great interest to observe fairly recent insertions from a large number of single-copy LTR retrotransposon families (**Supplementary Figure 22**).

The degree to which non-autonomous LTR retrotransposons impede the proliferation of autonomous retroelements has key evolutionary impact on the genome size (Zhang and Gao, 2017). We found a rapid and recent propagation of more than 4,000 nonautonomous elements (**Figure 2C**). Of them, some were derived from autonomous Ty3-*gypsy* or Ty1-*copia* families that have slowly lost internal protein-coding genes. However, it is difficult to determine counterpart autonomous families for others. The *TEL001* family was selected as an exemplar to show that partial and/or complete loss of internal protein-coding genes has resulted in a quick increase of incomplete autonomous and/or nonautonomous retroelements that have far exceeded autonomous ancestral elements within the last 1 Myr. Based on structural features of *TEL001*, 4,062 full-length LTR retrotransposons were classified into the four groups (**Figure 3B and 3C**; **Supplementary Figure 23**). Group 1 contained 451 copies with complete sequences of *gag* and *pol* (PR, RT and IN) genes; Group 2 comprised 352 copies with the loss of at least one of the *gag*, PR, RT and IN domains; Group 3 had 1,063 copies with only the *gag* domain; and Group 4 included 2,196 nonautonomous copies without any internal *gag* and *pol* genes. Due to the dominance of the nonautonomous elements, the proportion of effective retrotransposition-related source proteins possibly declined dramatically, and insertion rates of the entire *TEL001* family largely decreased most recently (**Figure 3B**). In addition to such nonautonomous elements derived from Ty3-*gypsy* or Ty1-*copia* source families, there were many nonautonomous families, such as *TEL004*, which is a very young family that has undergone a large number of recent insertions(**Supplementary Figure 21**). There were also many low-copy and single-copy nonautonomous families reproduced most recently, together making the recent inserted nonautonomous elements far exceed Ty3-*gypsy* or Ty1-*copia* copies (**Figure 2C**; **Supplementary Figure 22**; **Supplementary Figure 24**). We then assessed levels of gene expression of all types of LTR retrotransposons using Illumina RNA-seq data from the five tissues (**Figure 2D**; **Supplementary Tables 12 and 23**). We detected ∼16.70% (∼7,586) of all expressed transcripts and ∼10.38% of all mapped reads, on average, for five tissues that are associated with LTR retrotransposons (**Supplementary Table 23**). ∼63.59% of illumina reads mapped to multi-copy nonautonomous LTR retrotransposons families (e.g., *TEL004*, ∼45.88%; *TEL013* ∼7.03%; *TEL019*, ∼2.45%) exhibited notably high levels of gene expression than Ty1-*copia* and particularly Ty3-*gypsy* families in multi-copy families (**Figure 2D**; **Supplementary Table 23**). Proteins (including *gag*, PR, RT and IN domains in *pol*) necessary for the retrotransposition were further annotated using Pfam (Finn et al., 2013). Surprisingly, ∼94.23% of the expressed LTR retrotransposon-related transcripts were nothing related to encoding *gag* and *pol* genes and only 5.77% of the retrotransposon-related transcripts mapped to at least one of above-mentioned genes **(Supplementary Table 24)**. Our findings thus offer one more case that recently increased non-autonomous LTR retrotransposons with high expression levels may limit the efficiency by reducing the supply of enzymes needed for a successful retrotransposition (Zhang and Gao, 2017).

In conclusion, we have first generated a highly contiguous and accurate tea tree genome assembly of *C. sinensis* var. *sinensis* cv. *Biyun* using SMRT technology, which is much more improved compared to the formerly reported genome assembly of *C.sinensis* var. *sinensis* cv. *Shuchazao*. This effort has added one more successful example that sequencing the highly repetitive and heterozygous and relatively large tea tree genome may be achieved using high-depth long SMRT reads to resolve ambiguous genomic regions harboring predominantly repetitive sequences. Such a high-quality genome assembly of the tea tree is timely and will therefore be welcome to the broad tea research community, which is essential to enable researchers to accurately obtain functionally significant gene families that not only involve in the biosynthesis of numerous metabolites (e.g., catechins, theanine and caffeine) but also determine agronomically important traits relevant to the improvement of tea quality and production. The exceptionally contiguous and precise genome assembly of the tea tree is powerful to fully identify all types of long LTR retrotransposons and almost entirely characterize the abundance of retrotransposon diversity to resolve the nature of repetitive landscape of such a large genome. The evolutionary history of very recently augmented LTR retrotransposon families, which have not been done ever before, could now be tracked genome-wide by dating bursts of non-autonomous LTR retrotransposons and undertaking their interaction with autonomous LTR retrotransposons, afterwards driving the genome evolution.

## Methods

DNA was extracted from a *CSS-BY* individual collected in Yunnan Pu’er Tea Tree Breeding Station, Yunnan, China, for PacBio RSII and Hiseq X Ten sequencing platforms. We performed a PacBio-only assembly using an overlap layout-consensus method implemented in FALCON (version 0.3.0) (Chin et al., 2013). Considering the highly heterozygous nature of the tea tree genome, the pipeline of ‘Purge Haplotigs’ (Roach et al., 2018) was used to remove the redundant sequences caused by genomic heterozygosity. SSPACE-LongRead was subsequently employed to build scaffolds (English et al., 2012). The gaps were filled with all Pacbio subreads using PBJelly tool (English et al., 2012). Finally, we used Hi-C data to construct a high-quality chromosome-scale assembly using LACHESIS (Burton et al., 2013) and JUICERBOX (Durand et al., 2016; Robinson et al., 2018).

The Maker genome annotation pipeline (Cantarel et al., 2008) and five tissues transcriptome data (young leaf, YL; flower bud, FB; stem, ST; fruit, FR and tender shoot, TS) were used for gene prediction. The expression levels of annotated genes were computed by the pipeline of HISAT2 (V2.1.0) and StringTie (V1.3.5) (Pertea et al., 2016).

Five different types of non-coding RNA genes were predicted using various *de novo* and homology search methods. RepeatModeler (**Supplementary URLs**) was used to identify and model repeat families and statistic by RepeatMasker (version 4.0.5) (Chen, 2004; Smit et al., 2016) (**Supplementary URLs**). LTR_STRUCT (McCarthy and McDonald, 2003) was applied to identify LTR retrotransposon elements for the construction of *de novo* repeat library. SSRs were identified and located using MISA (**Supplementary URLs**).

Homologous genes from different plant species were combined using all *vs* all BLASTP (BLAST+ 2.71) (Altschul et al., 1997; Johnson et al., 2008) and then synteny blocks were identified with MCscanX (Wang et al., 2012). Gene families were clustered with OrthoMCL (Li et al., 2003).

## Accession Numbers

Raw PacBio and Illumina sequencing reads of *CSS-BY* have been deposited in the NCBI Sequence Read Archive Database under accession PRJNA381277. Genome assembly, gene prediction, gene functional annotations, and transcriptomic data may be accessed via the web site at: www.plantkingdomgdb.com/CSS-BY/.

## Supplementary Information

Supplementary Information is available at ***Molecular Plant*** Online.

## Author Contributions

L.-Z.G. designed and managed the project; C.S., Y.T., H.N., Y.-L.L., X.-L.Y. and X.-H. W. collected materials; C.S., C. L., C.-F.W. and X.-X.L. prepared and purified DNA and RNA samples; K.L. performed the genome assembly; Q.-J.Z., W.L., H.N., Y.Z., D.Z., W.-K.J. and Z.-Y.D. performed genome annotation and subsequent data analyses; L.-Z.G. and Q.-J.Z. wrote the manuscript; L.-Z.G., Q.-J. Z., Z.-H.L., X.-C.Z. and E.E.E. revised the manuscript.

## Acknowledgments

We appreciate the anonymous reviewers for their comments on this manuscript. The authors thank T. Brown for assistance in editing this manuscript. This study was supported by Startup Grant from South China Agricultural University and Yunnan Innovation Team Project (to L.Z. G.). E.E.E. is an investigator of the Howard Hughes Medical Institute.

## References

Altschul, S.F., Madden, T.L., Schäffer, A.A., Zhang, J., Zhang, Z., Miller, W., and Lipman, D.J. (1997). Gapped BLAST and PSI-BLAST: a new generation of protein database search programs. Nucleic acids research 25:3389–3402.

Ashburner, M., Ball, C.A., Blake, J.A., Botstein, D., Butler, H., Cherry, J.M., Davis, A.P., Dolinski, K., Dwight, S.S., and Eppig, J.T. (2000). Gene Ontology: tool for the unification of biology. Nature genetics 25:25.

Banerjee, B. (1992). Botanical classification of tea. In: Tea: Springer. 25–51.

Boeckmann, B., Bairoch, A., Apweiler, R., Blatter, M.-C., Estreicher, A., Gasteiger, E., Martin, M.J., Michoud, K., O’donovan, C., and Phan, I. (2003). The SWISS-PROT protein knowledgebase and its supplement TrEMBL in 2003. Nucleic acids research 31:365–370.

Burton, J.N., Adey, A., Patwardhan, R.P., Qiu, R., Kitzman, J.O., and Shendure, J. (2013). Chromosome-scale scaffolding of de novo genome assemblies based on chromatin interactions. Nature biotechnology 31:1119.

Cantarel, B.L., Korf, I., Robb, S.M., Parra, G., Ross, E., Moore, B., Holt, C., Alvarado, A.S., and Yandell, M. (2008). MAKER: an easy-to-use annotation pipeline designed for emerging model organism genomes. Genome research 18:188–196.

Chen, L., Zhou, Z.-X., and Yang, Y.-J. (2007). Genetic improvement and breeding of tea plant (Camellia sinensis) in China: from individual selection to hybridization and molecular breeding. Euphytica 154:239–248.

Chen, N. (2004). Using RepeatMasker to identify repetitive elements in genomic sequences. Current protocols in bioinformatics 5:4.10. 11-14.10. 14.

Chin, C.-S., Alexander, D.H., Marks, P., Klammer, A.A., Drake, J., Heiner, C., Clum, A., Copeland, A., Huddleston, J., and Eichler, E.E. (2013). Nonhybrid, finished microbial genome assemblies from long-read SMRT sequencing data. Nature methods 10:563.

Delcher, A.L., Salzberg, S.L., and Phillippy, A.M. (2003). Using MUMmer to identify similar regions in large sequence sets. Current protocols in bioinformatics:10.13. 11-10.13. 18.

Durand, N.C., Robinson, J.T., Shamim, M.S., Machol, I., Mesirov, J.P., Lander, E.S., and Aiden, E.L. (2016). Juicebox provides a visualization system for Hi-C contact maps with unlimited zoom. Cell systems 3:99–101.

English, A.C., Richards, S., Han, Y., Wang, M., Vee, V., Qu, J., Qin, X., Muzny, D.M., Reid, J.G., and Worley, K.C. (2012). Mind the gap: upgrading genomes with Pacific Biosciences RS long-read sequencing technology. PloS one 7:e47768.

Finn, R.D., Bateman, A., Clements, J., Coggill, P., Eberhardt, R.Y., Eddy, S.R., Heger, A., Hetherington, K., Holm, L., and Mistry, J. (2013). Pfam: the protein families database. Nucleic acids research 42:D222–D230.

Hřibová, E., Neumann, P., Matsumoto, T., Roux, N., Macas, J., and Doležel, J. (2010). Repetitive part of the banana (*Musa acuminata*) genome investigated by low-depth 454 sequencing. BMC plant biology 10:204.

Huang, S., Ding, J., Deng, D., Tang, W., Sun, H., Liu, D., Zhang, L., Niu, X., Zhang, X., and Meng, M. (2013). Draft genome of the kiwifruit Actinidia chinensis. Nature communications 4:2640.

Johnson, M., Zaretskaya, I., Raytselis, Y., Merezhuk, Y., McGinnis, S., and Madden, T.L. (2008). NCBI BLAST: a better web interface. Nucleic acids research 36:W5–W9.

Jones, P., Binns, D., Chang, H.-Y., Fraser, M., Li, W., McAnulla, C., McWilliam, H., Maslen, J., Mitchell, A., and Nuka, G. (2014). InterProScan 5: genome-scale protein function classification. Bioinformatics 30:1236–1240.

Kanehisa, M., and Goto, S. (2000). KEGG: kyoto encyclopedia of genes and genomes. Nucleic acids research 28:27–30.

Li, C.-F., Zhu, Y., Yu, Y., Zhao, Q.-Y., Wang, S.-J., Wang, X.-C., Yao, M.-Z., Luo, D., Li, X., and Chen, L. (2015). Global transcriptome and gene regulation network for secondary metabolite biosynthesis of tea plant (Camellia sinensis). BMC genomics 16:560.

Li, L., Stoeckert, C.J., and Roos, D.S. (2003). OrthoMCL: identification of ortholog groups for eukaryotic genomes. Genome research 13:2178–2189.

Li, S., Lo, C.-Y., Pan, M.-H., Lai, C.-S., and Ho, C.-T. (2013). Black tea: chemical analysis and stability. Food & function 4:10–18.

Liu, Z., Gao, L., Chen, Z., Zeng, X., Huang, J.a., Gong, Y., Li, Q., Liu, S., Lin, Y., Cai, S., et al. (2019). Leading progress on genomics, health benefits and utilization of tea resources in China. Nature.

Llorens, C., Munoz-Pomer, A., Bernad, L., Botella, H., and Moya, A. (2009). Network dynamics of eukaryotic LTR retroelements beyond phylogenetic trees. Biol Direct 4:41.

McCarthy, E.M., and McDonald, J.F. (2003). LTR_STRUC: a novel search and identification program for LTR retrotransposons. Bioinformatics 19:362–367.

Ming, T., and Bartholomew, B. (2007). Theaceae. In Flora of China, Z. Wu, P. Raven, and D. Hong, eds. (Beijing and St. Louis: Science Press and Missouri Botanical Garden):pp. 367–412.

Möller, S., Croning, M.D., and Apweiler, R. (2001). Evaluation of methods for the prediction of membrane spanning regions. Bioinformatics 17:646–653.

Mondal, T.K., Bhattacharya, A., Laxmikumaran, M., and Ahuja, P.S. (2004). Recent advances of tea (Camellia sinensis) biotechnology. Plant Cell, Tissue and Organ Culture 76:195–254.

Narukawa, M., Morita, K., and Hayashi, Y. (2008). L-theanine elicits an umami taste with inosine 5′-monophosphate. Bioscience, biotechnology, and biochemistry 72:3015–3017.

Parra, G., Bradnam, K., and Korf, I. (2007). CEGMA: a pipeline to accurately annotate core genes in eukaryotic genomes. Bioinformatics 23:1061–1067.

Pertea, M., Kim, D., Pertea, G.M., Leek, J.T., and Salzberg, S.L. (2016). Transcript-level expression analysis of RNA-seq experiments with HISAT, StringTie and Ballgown. Nature protocols 11:1650.

Piegu, B., Guyot, R., Picault, N., Roulin, A., Sanyal, A., Kim, H., Collura, K., Brar, D.S., Jackson, S., Wing, R.A., et al. (2006). Doubling genome size without polyploidization: dynamics of retrotransposition-driven genomic expansions in *Oryza australiensis*, a wild relative of rice. Genome research 16:1262–1269.

Roach, M.J., Schmidt, S.A., and Borneman, A.R. (2018). Purge Haplotigs: allelic contig reassignment for third-gen diploid genome assemblies. BMC bioinformatics 19:460.

Robinson, J.T., Turner, D., Durand, N.C., Thorvaldsdottir, H., Mesirov, J.P., and Aiden, E.L. (2018). Juicebox. js provides a cloud-based visualization system for Hi-C data. Cell systems 6:256-258. e251.

Shi, C.-Y., Yang, H., Wei, C.-L., Yu, O., Zhang, Z.-Z., Jiang, C.-J., Sun, J., Li, Y.-Y., Chen, Q., and Xia, T. (2011). Deep sequencing of the Camellia sinensis transcriptome revealed candidate genes for major metabolic pathways of tea-specific compounds. BMC genomics 12:131.

Shi, T., Huang, H., and Barker, M.S. (2010). Ancient genome duplications during the evolution of kiwifruit (Actinidia) and related Ericales. Annals of Botany 106:497–504.

Simão, F.A., Waterhouse, R.M., Ioannidis, P., Kriventseva, E.V., and Zdobnov, E.M. (2015). BUSCO: assessing genome assembly and annotation completeness with single-copy orthologs. Bioinformatics 31:3210–3212.

Smit, A., Hubley, R., and Green, P. (2016). RepeatMasker Open-4.0. 2015. Google Scholar.

Soni, R.P., Katoch, M., Kumar, A., Ladohiya, R., and Verma, P. (2015). Tea: Production, composition, consumption and its potential an antioxidant and antimicrobial agent. International Journal of Food and Fermentation Technology 5:95.

Taniguchi, F., Kimura, K., Saba, T., Ogino, A., Yamaguchi, S., and Tanaka, J. (2014). Worldwide core collections of tea (Camellia sinensis) based on SSR markers. Tree genetics & genomes 10:1555–1565.

Vitte, C., and Panaud, O. (2005). LTR retrotransposons and flowering plant genome size: emergence of the increase/decrease model. Cytogenet Genome Res 110:91–107.

Vitte, C., Panaud, O., and Quesneville, H. (2007). LTR retrotransposons in rice (*Oryza sativa*, L.): recent burst amplifications followed by rapid DNA loss. BMC Genomics 8:218.

Wang, Y., Tang, H., DeBarry, J.D., Tan, X., Li, J., Wang, X., Lee, T.-h., Jin, H., Marler, B., and Guo, H. (2012). MCScanX: a toolkit for detection and evolutionary analysis of gene synteny and collinearity. Nucleic acids research 40:e49–e49.

Wei, C., Yang, H., Wang, S., Zhao, J., Liu, C., Gao, L., Xia, E., Lu, Y., Tai, Y., and She, G. (2018). Draft genome sequence of Camellia sinensis var. sinensis provides insights into the evolution of the tea genome and tea quality. Proceedings of the National Academy of Sciences:201719622.

Wheeler, D.S., and Wheeler, W.J. (2004). The medicinal chemistry of tea. Drug development research 61:45–65.

Wicker, T., and Keller, B. (2007). Genome-wide comparative analysis of *copia* retrotransposons in Triticeae, rice, and *Arabidopsis* reveals conserved ancient evolutionary lineages and distinct dynamics of individual *copia* families. Genome research 17:1072–1081.

Willson, K.C., and Clifford, M.N. (2012). Tea: Cultivation to consumption: Springer Science & Business Media.

Xia, E.-H., Zhang, H.-B., Sheng, J., Li, K., Zhang, Q.-J., Kim, C., Zhang, Y., Liu, Y., Zhu, T., and Li, W. (2017). The tea tree genome provides insights into tea flavor and independent evolution of caffeine biosynthesis. Molecular plant 10:866–877.

Zhang, Q.-J., and Gao, L.-Z. (2017). Rapid and recent evolution of LTR retrotransposons drives rice genome evolution during the speciation of AA-genome Oryza species. G3: Genes, Genomes, Genetics:g3. 116.037572.

