## Supplementary Information and Figures for "SMRT sequencing yields the chromosome-scale reference genome of tea tree, *Camellia sinensis* var. *sinensis*"

### Supplementary Methods

#### Plant Materials, Genome Sequencing, and Assembly

Fresh and healthy leaves were harvested from an individual plant of cultivar *Camellia sinensis* cv. *Biyun* (*CSS-BY*) from Pu'er City, Yunnan Province, China. They were immediately frozen in liquid nitrogen after collection, followed by the preservation at -80°C in the laboratory prior to DNA extraction. Total DNA was extracted for sequencing on the PacBio RSII and HiSeq X Ten sequencing platforms. ~282.94 Gb of Illumina sequencing data with 250 bp insert size on the HiSeq X Ten platform **(Supplementary Table 1)** were generated to polish the genome assembly using Pilon (Walker et al., 2014). The SMRTbell template libraries were prepared following the standard protocol for long-insert libraries according to the manufacturer's instructions (Pacific Biosciences of California, Inc. or PacBio). Genomic DNA was firstly sheared to a size range of 15-40 kb using a Covaris® g-TUBE device (Covaris). The large fragments were enriched and enzymatically repaired and converted into the three SMRTbell template libraries. These fragments were ligated with hairpin adapters. The remaining damaged DNA fragments and those without adapters at both ends were eliminated by digestion with exonucleases. The resulting SMRTbell templates were size-selected from 7 kb to 50 kb by Blue Pippin electrophoresis (Sage Sciences) and sequenced on a Sequel instrument using P6-C4 sequencing chemistry. Data were collected as 10 h sequencing videos and 1,679.6 Gb of raw sequencing data were generated **(Supplementary Table 2)**.

For Hi-C sequencing, chromosomal structure was fixed by formaldehyde crosslinking, and then *MboI* enzyme was used to shear DNA. Hi-C library with 200-600 bp insert size was constructed, which was sequenced on Hiseq 2000 platform. The Hi-C sequence data were qualified with HIC-pro (Servant et al., 2015), in which the validly mapped reads were selected to cluster and order the assembled genome sequences by the ALLHiC pipeline (Dudchenko et al., 2017) **(Supplementary Table 3)**.

#### Genome Assembly and Quality Control

In order to estimate the genome size of *C. sinensis* var. *sinensis* cv. *Biyun* (*CSS-BY*), high-quality Illumina short-insert reads were used to examine the 17-mer sequences using sliding windows. The frequency of occurrence distribution of 17-mer sequences were calculated (**Supplementary Figure 1).** The genome size was estimated based on the formula 'Genome size = total *K*-mer num / *K*-mer depth'. The main volume peak of *K*-mer frequency was 69, so the genome size was estimated as ~3.25 Gb based on this formula. The peak before the main peak showed that the genome was a highly heterozygous genome.

*De novo* assembly of PacBio SMRT reads was performed using FALCON software (3.1). All reads were first pairwise aligned, and the top 60 long coverage of subreads were selected as seeds to perform error correction with parameters '--output_multi --min_idt 0.70 --min_cov 4 --max_n_read 300'. About 36.6 × coverage of corrected reads with N50 length of 14.6 kb were remained and aligned to each other to construct string graphs. Graph were further filtered with parameters '--max_diff 70 --max_cov 70 --min_cov 3' to generate contigs. After the initial assembly, the produced contigs were then polished by aligning SMRT reads using Arrow from the SMRTlink 5.0 software (PacBio) with default parameters. Then, Pilon (Walker et al., 2014) was used to perform the second round of error correction with the Illumina paired-end reads. The total length of the assembly was ~4.41 Gb with contig N50 of ~403.63 Kb (V1.0 in **Supplementary Table 4)**. The 'Purge Haplotigs' pipeline (Roach et al., 2018) was used to remove the redundant sequences caused by heterozygosity. After removing the redundant sequences, 11,100 contigs were remained with a total length of ~3.08 Gb and the contig N50 size was improved to ~562.48 Kb (V1.1 in **Supplementary Table 4)**. To further improve the continuity of the assembly, SSPACE-LongRead was used to build scaffolds. The PBJelly tool (English et al., 2012) was used to fill gaps with all PacBio subreads, generating a genome with a contig N50 size of ~671.89 Kb and scaffold N50 size of ~1.09 Mb. The total length of this version

(V1.2 in **Supplementary Table 4)** was ~3.13 Gb, containing 0.14% Ns. The 'Purge Haplotigs' was used to remove the redundant sequences once again and the final assembly length was ~2.92 Gb with contig N50 size of ~707.72 Kb and scaffold N50 length of ~1.16 Mb (V1.3 in **Supplementary Table 4)**.

For chromosome-level assembly, the high-quality Hi-C data were mapped to the original scaffolds using BWA v0.7.7 (ref. 11), and broke the assembly error by abnormal coverage. Only reads with unique alignment positions were extracted to construct a chromosome-scale assembly using LACHESIS (Burton et al., 2013) and JUICERBOX(Durand et al., 2016; Robinson et al., 2018). In total, 6,590 contigs (representing ~98.45% of the total genome sequencing length) were anchored to the 15 chromosomes of *CSS-BY*. The contact maps of each chromosome were visualized with heatmaps at **Supplementary Figure 2**. The final assembly had the GC content of 38.24% with the longest contig and scaffold at ~3.91 Mb and ~238.63 Mb, respectively **(Supplementary Tables 4-6)**.

To evaluate the completeness of the tea tree genome assembly, we applied both CEGMA (Core Eukaryotic Gene Mapping Approach) (Parra et al., 2007) and BUSCO (Benchmarking Universal Single-Copy Orthologs) (Simão et al., 2015) to evaluate the assembly completeness. We also estimated the mapping rate and coverage of genome assembly by mapping the Illumina paired-end reads to our assembly using BWA (Li and Durbin, 2009).

**Genome Annotation**

We used the Maker genome annotation pipeline for gene prediction (Cantarel et al., 2008). Gene evidence was provided by 467,320 transcripts, which were generated by assembling RNA-Seq reads from young leaf (YL), flower bud (FB), stem (ST), fruit (FR) and tender shoot (TS) using Trinity (version r2.02) (**Supplementary Tables 12-13**) (Grabherr et al., 2011). The final gene set was generated after removing weakly supported gene models by following filtering thresholds: (1) No TE, mRNA length >=

200 bp, CDS length >= 30 bp; (2) if 0.1 < AED <= 0.4, FPKM must higher than 0.5; (3) if AED > 0.4, FPKM must higher than 0.5 and CDS need Pfam domain support. For function annotation, amino acid sequences were searched against known genes of SwissProt (Boeckmann et al., 2003), Pfam (Finn et al., 2013), TmHMM (Möller et al., 2001), KEGG (Kanehisa and Goto, 2000)， GO (Ashburner et al., 2000) and InterProScan (Jones et al., 2014) databases (**Supplementary Table 14; Supplementary Figure 5-6**).

Five different types of non-coding RNA genes were predicted **(Supplementary Table 15)**. The tRNA genes were identified by the tRNAscan-SE algorithms (version 1.23) (Lowe and Eddy, 1997); the rRNA genes (8S, 18S, and 28S) were predicted using the RNAmmer algorithms (Lagesen et al., 2007); the snoRNA genes were annotated using snoScan (Lowe and Eddy, 1999); the snRNA genes were identified by INFERNAL software against the Rfam database (release 9.1) (Griffiths-Jones et al., 2005; Nawrocki et al., 2009); for miRNA genes, the conserved miRNAs were identified by mapping all miRBase-recorded plant miRNA precursor sequences (Kozomara and Griffiths-Jones 2011) against the assembled *CSS-BY* genome using BLASTN (BLAST+ 2.71) (Altschul et al., 1997; Johnson et al., 2008). When a miRNA was mapped to a target *CSS-BY* genome, the surrounding sequences were then checked for hairpin structures. Those loci that fulfilled miRNA precursor secondary structures were annotated as miRNA genes.

**Gene Expression Analysis**

The RNA_Seq data from five tissues (YL, TS, FB, FR and ST) were sequenced on HiSeq X Ten platform, and ~26.22 Gb of Illumina sequencing data was generated (**Supplementary Table 12**). The clean data were mapped with HISAT2 (V2.1.0) to the annotated genome, and then the transcripts were next determined by StringTie (V1.3.5) (Pertea et al., 2016). The gene expression levels were computed as the number of reads per kilobase of gene length per million mapped reads (FPKM).

**Analysis of Repetitive Elements**

Repetitive elements were identified based on homologous detection and *de novo* searches. RepeatModeler (**Supplementary URLs**) was used to identify and model repeat families. Then, RepeatMasker (version 4.0.5) (Chen, 2004; Smit et al., 2016) (**Supplementary URLs**) was used to annotate and mask repetitive elements using the library generated by RepeatModeler. Simple Sequence Repeats (SSRs) were identified and located using MISA (**Supplementary URLs**). LTR_STRUCT (McCarthy and McDonald, 2003) was applied to identify LTR retrotransposons for the construction of a *de novo* repeat library, and genomic locations were also detected using RepeatMasker (v4.0.6) (Chen, 2004; Smit et al., 2016). All full-lengthLTR retrotransposons (the full-length retroelements here refer to those LTR-RTs which had the structure of two LTRs, PBS and PPT, which can define a repeat element to LTR-RT ) were classified into Ty1-*copia*, Ty3-*gypsy* and unclassified groups according to the order of ORFs using Pfam (Finn et al., 2008). To compare the structure and encoding potential, we divide LTR-RTs into three categories: complete autonomous elements (Group 1 in **Figure 3B**) which include all retro-transposition needed protein, incomplete autonomous elements (Group 2 & 3 in **Figure 3B**) which include part of retro-transposition needed protein, and non-autonomous elements (Group 4 in **Figure 3B**) which do not include retrotransposition needed proteins.

Amino acid sequence alignments for LTR retrotransposons were performed using Clustal W2 (Larkin et al., 2007). While constructing the phylogenetic tree, we removed sequences with premature stop codons . The neighbor joining (NJ) method was used to generate unrooted trees using uncorrected pairwise distances from the sequence alignments with the program PHYLIP (v3.697) (Felsenstein, 1993). ~151 aa for Ty3-*gypsy* and ~93 aa for Ty1-*copia* were used to build NJ trees. Then, LTR retrotransposons were classified into families using BLASTClust (in v2.2.11 BLAST package) (**Supplementary URLs**) and all-to-all BLAST (v2.71 BLAST+) (Altschul et al., 1997; Johnson et al., 2008) of 5' LTR / 3' LTR sequences, followed by manual inspection. The family classification standard was considered acceptable if more than 80% of the 5' LTRs / 3' LTRs and sequence identity were greater than 80% (Llorens et al., 2011; Wicker et al., 2007). The complete autonomous, incomplete autonomous and non-autonomous LTR retrotransposons with similar LTRs were annotated and usually classified into the same family. Only when a LTR retrotransposon family had no autonomous counterpart, we define it as a non-autonomous LTR retrotransposon family.

The insertion times of LTR retrotransposons were calculated based on a previously published approach (SanMiguel et al., 1998). The two LTRs of each LTR retrotransposon that contains target-site duplication (TSD) were aligned using ClustalW (Larkin et al., 2007), and their nucleotide divergence was estimated using the baseml module implemented in PAML (Yang, 2007). The insertion times were then computed using: $T$ = K/2r; where: $T$ = insertion time; r = synonymous mutations/site/Myr; K = the divergence between the two LTRs (Baucom et al., 2009; Vitte et al., 2007). As the substitution rate of LTR retrotransposons was believed to be approximately twofold higher than that determined by the coding regions (Ma and Bennetzen 2004), we employed an average substitution rate of (r) $5.62 \times 10^{-9}$ substitutions per synonymous site per year to estimate insertion times of LTR retrotransposons (Huang et al., 2013; Shi et al., 2010). HISAT2 (V2.1.0) and StringTie (V1.3.5) (Pertea et al., 2016) were used to assemble the transcripts without a GTF file. Next, all the obtained transcripts were align to LTR retrotransposons by BLAST+ (v2.71) (Altschul et al., 1997; Johnson et al., 2008) to check their family classification. Then Pfam (Finn et al., 2013) was used to check the expressed *gag* and *pol* (PR, RT and IN) genes of LTR retrotransposons. HTSeq (v0.6.1p1) (Anders et al., 2015) was used to count reads mapped to LTR retrotransposon transcripts.

**Genome Collinearity and Gene Family Annotation**

Homologous genes from different plant species were combined using all *vs* all BLASTP (BLAST+ 2.71) (Altschul et al., 1997; Johnson et al., 2008). Gene families were clustered with OrthoMCL (Li et al., 2003) according to BLAST results.

Homologous genes were detected by BLASTP (BLAST+ 2.71) (Altschul et al., 1997; Johnson et al., 2008), and then synteny blocks were identified with MCscanX (Wang et al., 2012). The genome collinearity between the two assembled *CSS* genomes were performed by MCscanX (Wang et al., 2012) and Mummer 4 (Delcher et al., 2003), and the graph was drawn by TBtools (Chen et al., 2018).

To identify key genes involved in the three important metabolic pathways (the biosynthesis of catechins, theanine and caffeine) in *CSS-BY*, we used BLASTP (BLAST+ 2.71) (Altschul et al., 1997; Johnson et al., 2008) to find relevant proteins based on amino acid sequences of key genes previously reported in *CSA-YK10* (Xia et al., 2017), *CSS-SCZ* (Wei et al., 2018), *Actinidia chinensis* (kiwifruit) (Yao et al., 2015) and *Arabidopsis thaliana* (NCBI). Then, the matched genes were further curated using NR databases and incorrect proteins were manually removed. Multiple sequence alignments were performed using ClustalW2 (Larkin et al., 2007); and the maximum likelihood (ML) trees with the JTT model and 1000 bootstrap replicates were built using MEGA (version 7.0.18) (Tamura et al., 2007).

#### Supplementary URLs

BLASTClust: http://www.ncbi.nlm.nih.gov/Web/Newsltr/Spring04/blastlab.html;

MISA Perl script: http://pgrc.ipk-gatersleben.de/misa/;

Pfam database: http://pfam.xfam.org/;

psRNATarget: http://plantgrn.noble.org/psRNATarget/;

RepeatMasker: http://repeatmasker.org/;

RepeatModeler: http://www.repeatmasker.org/RepeatModeler.html;

The R project for statistical computing: https://www.r-project.org/.

### Supplementary Figures

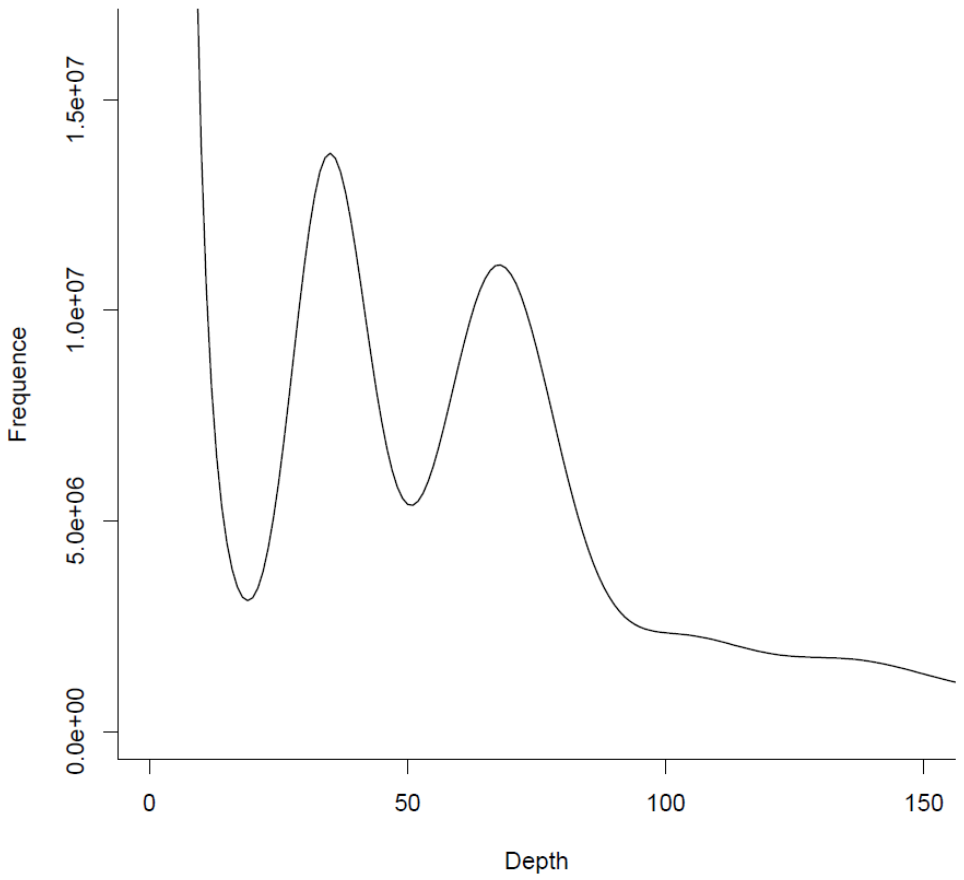


**Supplementary Figure 1. The 17-*mer* distribution of sequencing reads from the *C. sinensis* var. *sinensis* cv. *Biyun* (*CSS-BY*) genome**. The 17-*mer* occurrence was calculated using jellyfish (version 2.1.3) based on sequencing data from short insert size libraries (insert size ≤ 500 bp) of the tea tree genome. The main peak of K-*mer* frequency was 69, and thus the genome size was estimated at 3.25 Gb. Notably, the sharp peak before the main peak indicates that the genome is highly heterozygous.

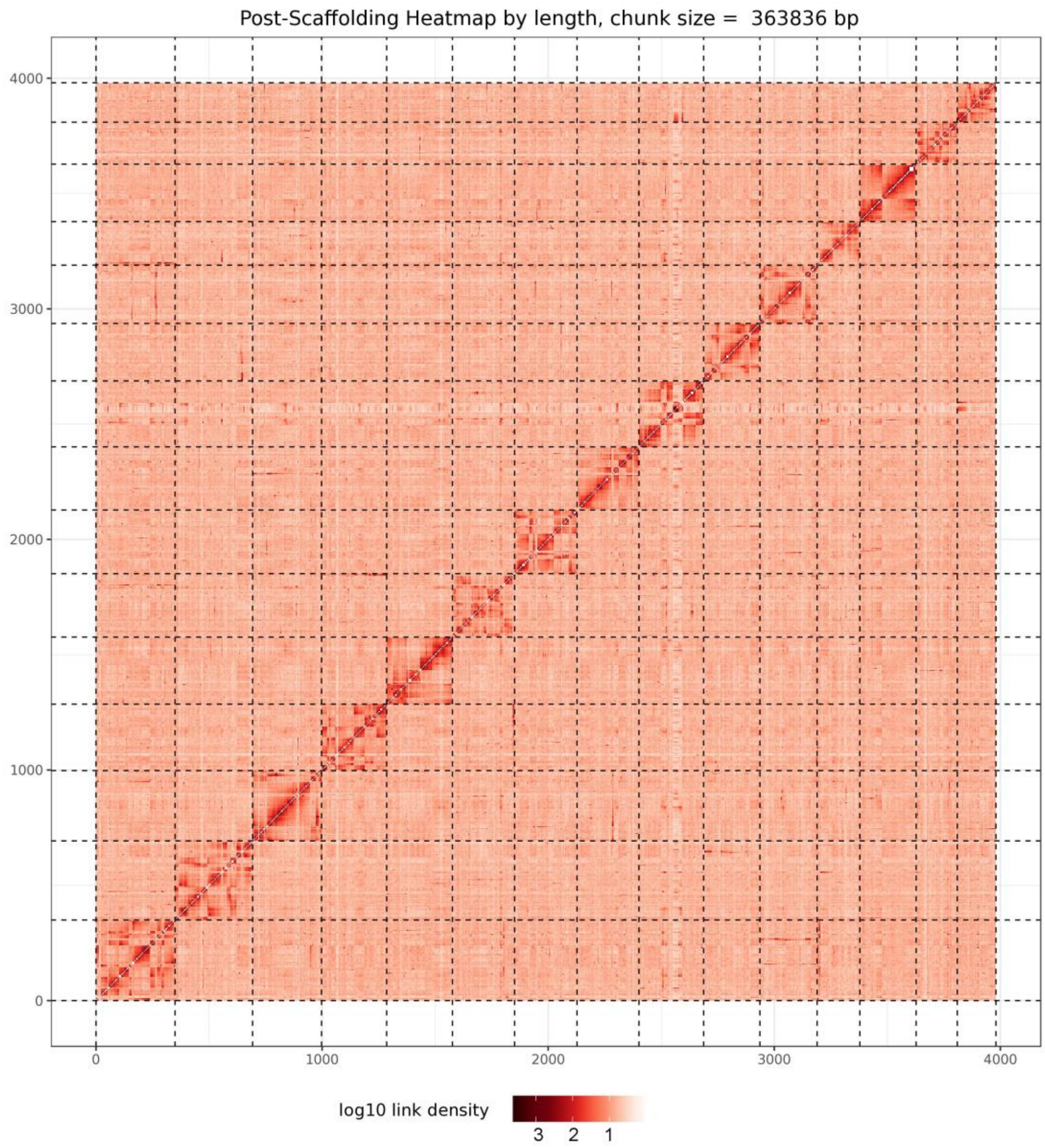


**Supplementary Figure 2. Hi-C clustering for pseudomolecule construction in *C. sinensis* var. *sinensis* cv. *Biyun*.** Post-clustering heat map of HiC-based intrachromosomal interactions in *C. sinensis* var. *sinensis* cv. *Biyun*. Pseudomolecules corresponding to the 15 haploid chromosomes are delineated by gray boxes.

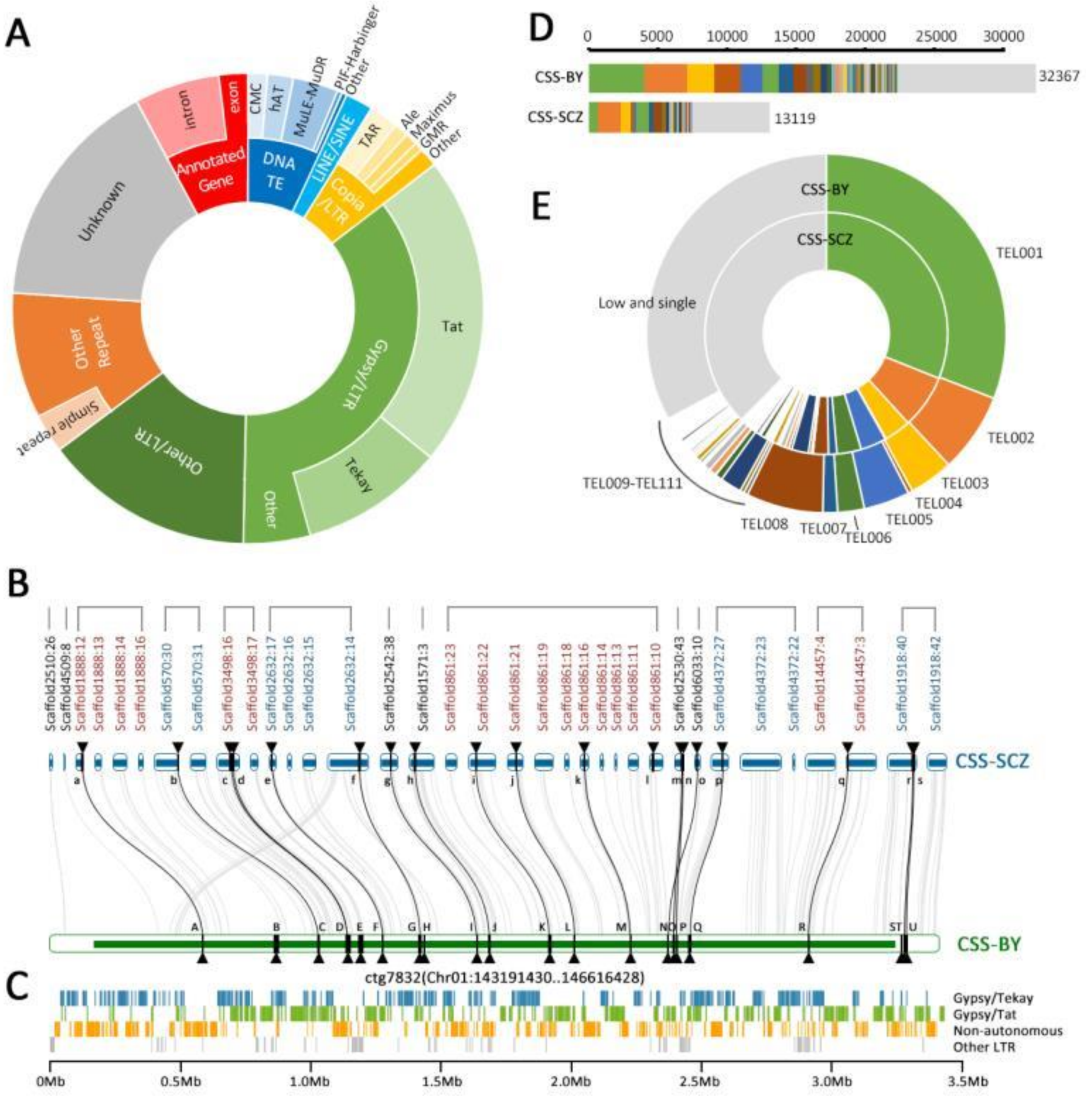


**Supplementary Figure 3. Genome components of *C. sinensis* var. *sinensis* cv. *Biyun*.** (**A**) Genome constituents of annotated genes and repeat sequences of *CSS-BY*. (**B**) Genomic region comparisons between the two assembled tea tree genomes, *CSS-BY* and *CSS-SCZ*, at the contig level. The ctg7832:1 from *CSS-BY* is corresponding to 35 contigs (derived from 14 scaffolds) in the *CSS-SCZ* genome assembly. The ctg7832:1 stands for the scaffold name, and the number after ':' denotes contig number in the scaffold; the same rule is adopted for the nomination of the *CSS-SCZ* contigs. The black triangles signify the annotated genes: a for *TEA016102*, b for *TEA012981*, c for *TEA032615*, d for *TEA032609*, e for *TEA024516*, f for *TEA024507*, g for *TEA000014*, h for *TEA031065*, i for *TEA018572*, j for *TEA018584*, k for *TEA018594*, l for *TEA031863*, m for *TEA031858*, n for *TEA018576*, o for *TEA021888*, p for *TEA014907*, q for *TEA011378*, r for *TEA021564*, and s for *TEA021563* in CSS-SCZ; A for *CSS00089*, B for *CSS00089*, C for *CSS00089*, D for *CSS00089*, E for *CSS00089*, F for *CSS00089*, G for *CSS00089*, H for *CSS00089*, I for *CSS00089*, J for *CSS00089*, K for *CSS00090*, L for *CSS00090*, M for *CSS00090*, N for *CSS00090*, O for *CSS00090*, P for *CSS00090*, Q for *CSS00090*, R for *CSS00090*, S for *CSS00090*, T for *CSS00090*, U for *CSS00091*, V for *CSS00091*, W for *CSS00091*, X for *CSS00091*, Y for *CSS00091* and Z for *CSS00091* in CSS-BY. Black lines indicate the gene collinearity, and gray lines show the synteny of intergenic regions. (**C**) Genomic distribution of LTR retrotransposons in ctg7832_pilion:1 of *CSS-BY*. (**D**) Comparisons of copy number of full-length LTR retrotransposon families between the two assembled tea tree genomes of *C. sinensis* var. *sinensis*. (**E**) Sequence length comparisons of LTR retrotransposon families annotated in the two assembled tea tree genomes of *C. sinensis* var. *sinensis*.

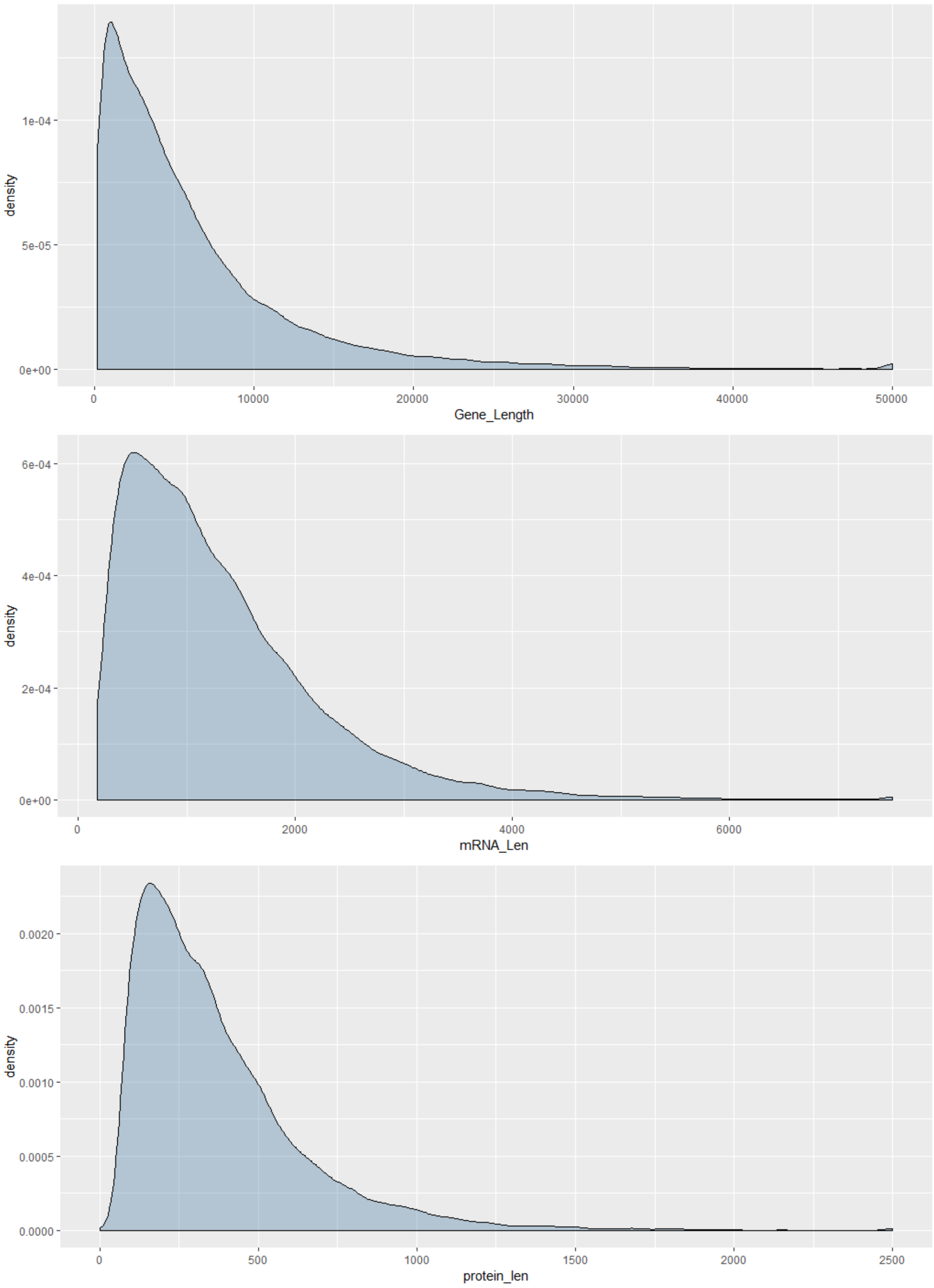


**Supplementary Figure 4. Statistics for gene, mRNA and protein lengths of the *CSS-BY* genome.**

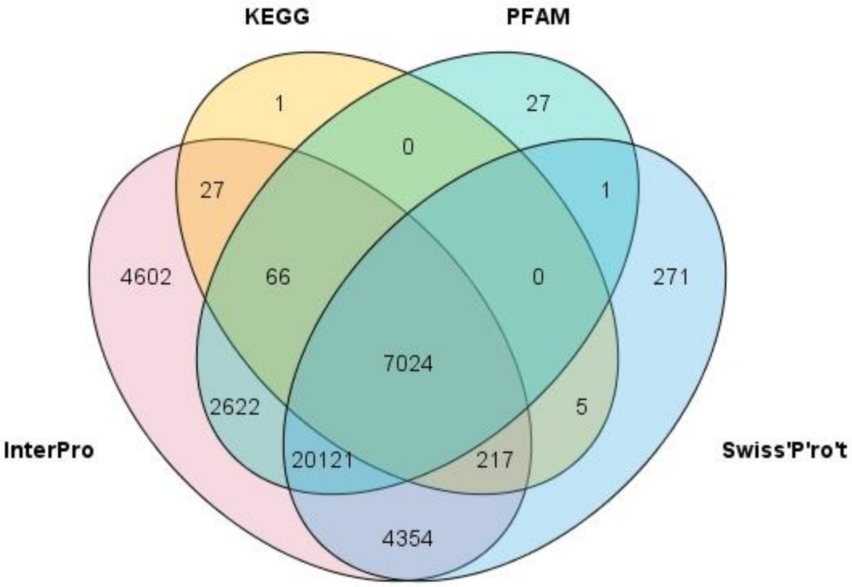


**Supplementary Figure 5. Venn diagram of gene function of the *CSS-BY* genome.**

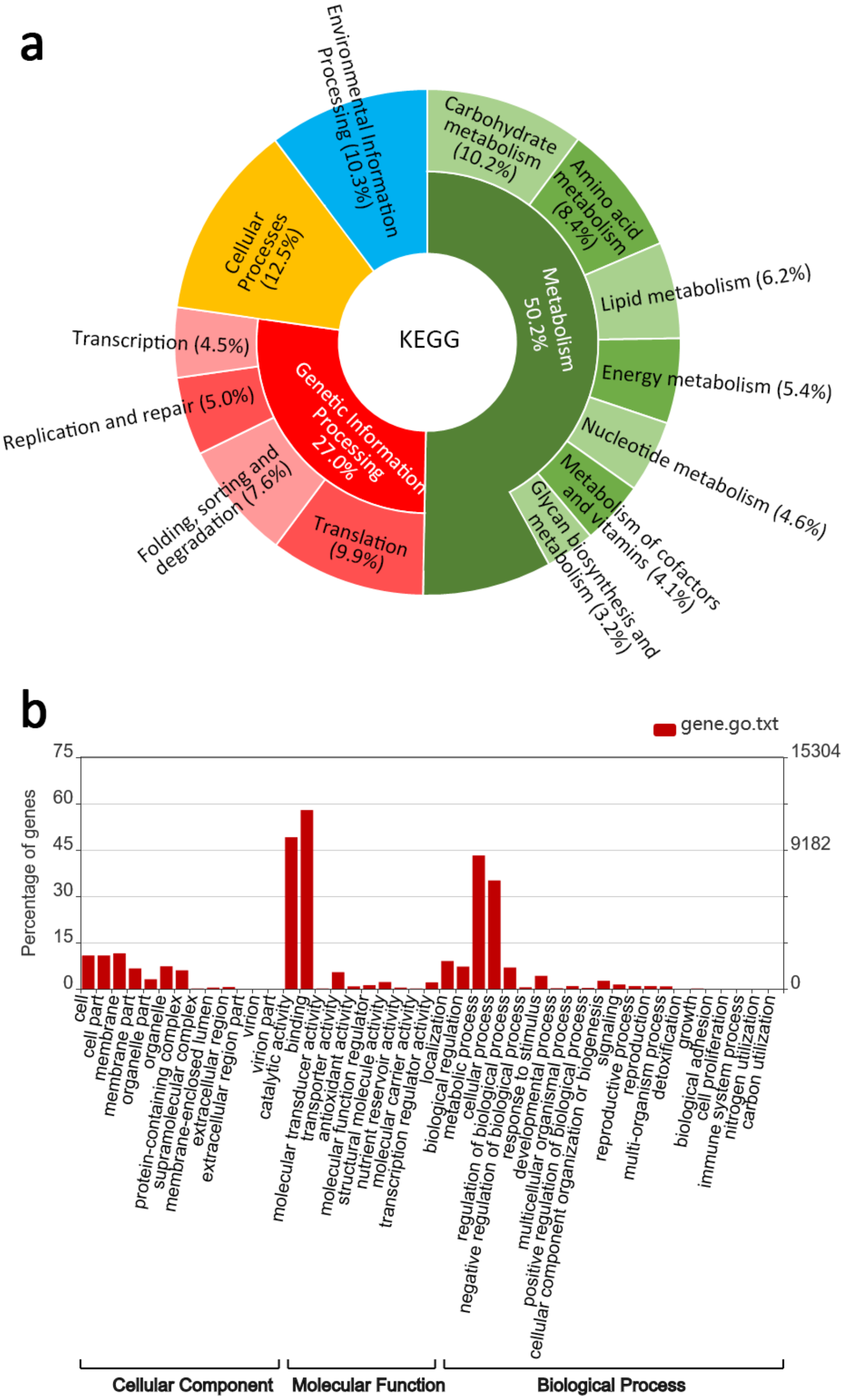


**Supplementary Figure 6. Statistic gene annotation in the *CSS-BY* genome. (a)** Distribution of predicted genes among levels 1 and 2 of Kyoto Encyclopedia of Genes and Genomes (KEGG) pathway. **(b)** Distribution of predicted genes among high-level Gene Ontology (GO) biological process terms.

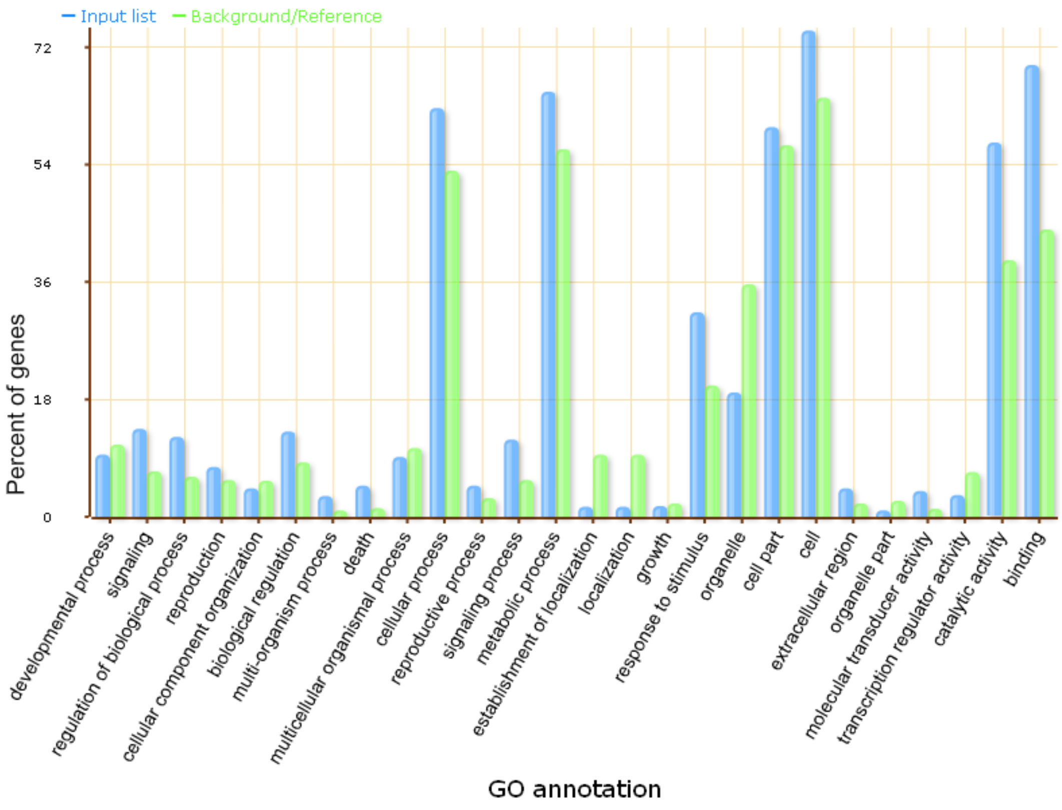


**Supplementary Figure 7. GO annotation of miRNA target genes in the *CSS-BY* genome.**

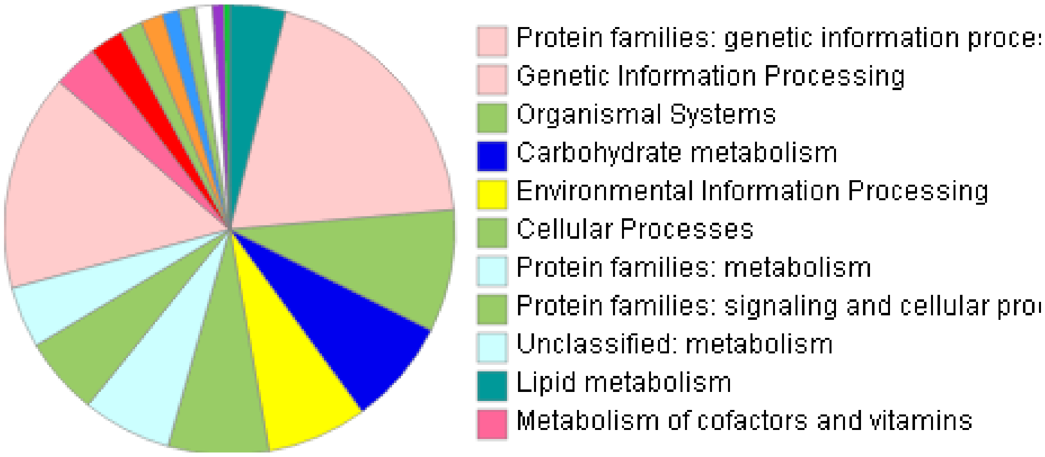


**Supplementary Figure 8. KO annotation of miRNA target genes in the *CSS-BY* genome.**

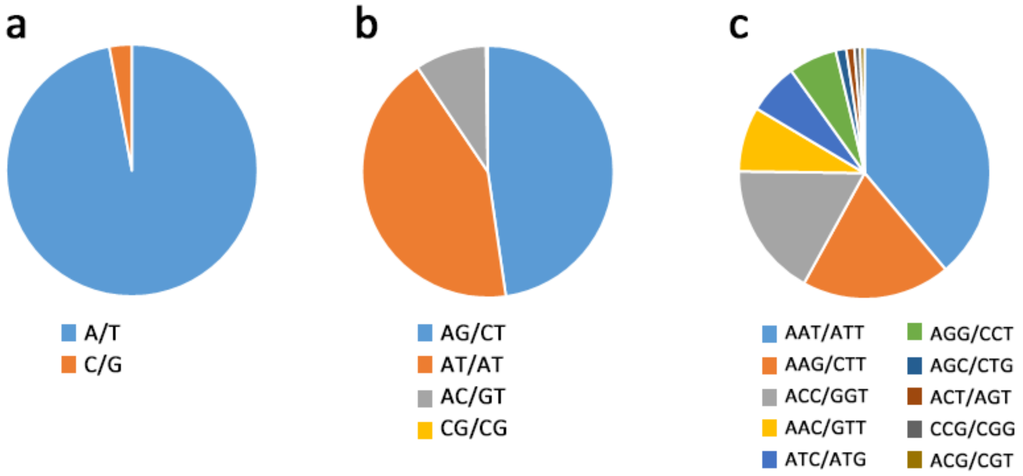


**Supplementary Figure 9. Percentages of different types of simple sequence repeats in the *CSS-BY* genome. (a)** Monomer. **(b)** Dimer. **(c)** Trimmer.

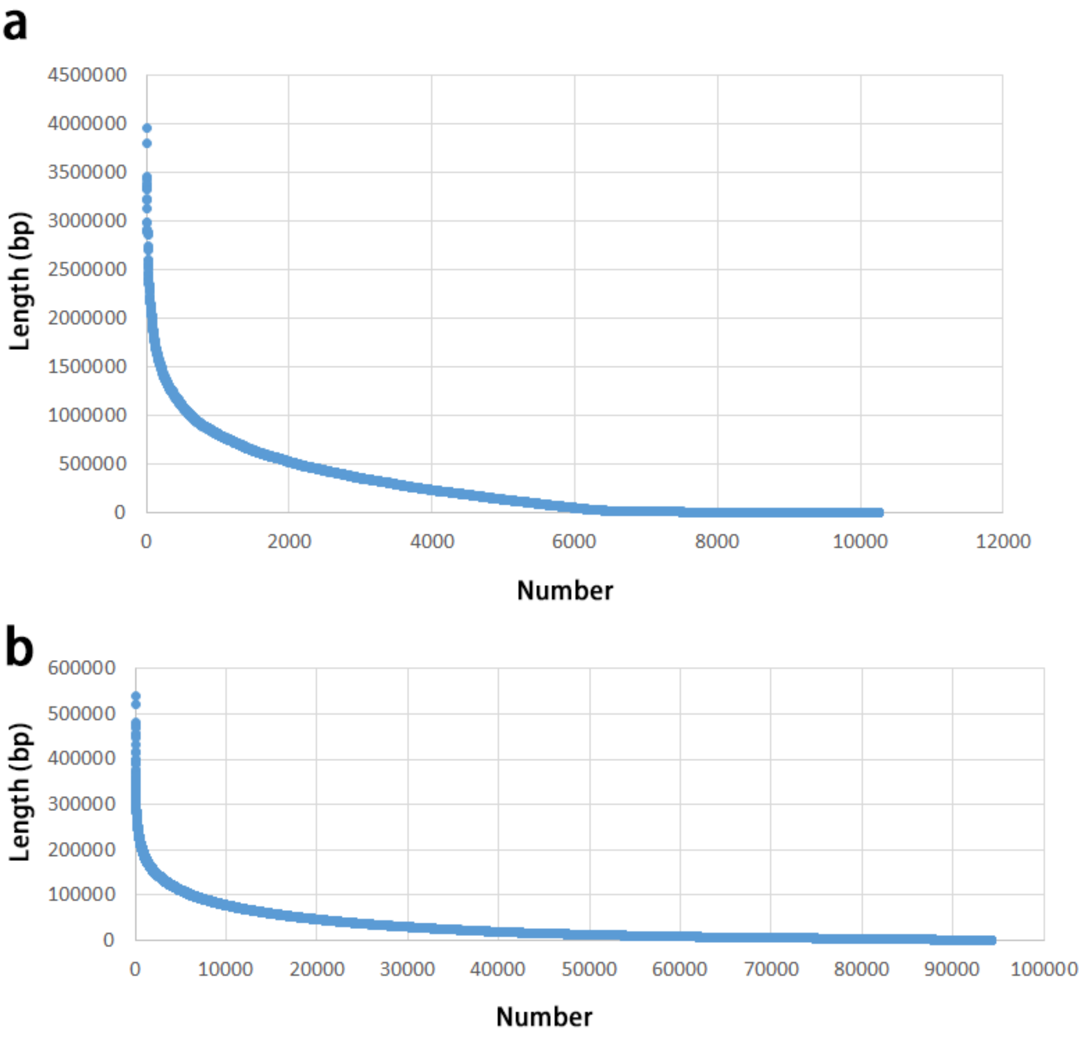


**Supplementary Figure 10. Contig length comparison of the two genome assemblies of *C. sinensis* var. *sinensis*: (a)** *CSS-BY*, and **(b)** *CSS-SCZ*.

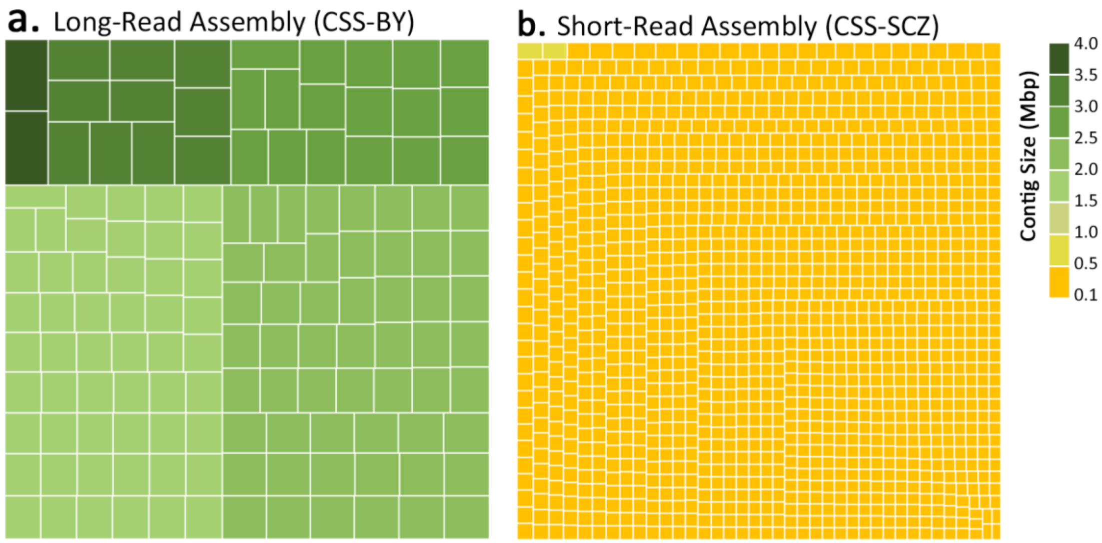


**Supplementary Figure 11. Tree maps representing differences in the fragmentation of the long-read and short-read genome assemblies of *C. sinensis* var. *sinensis*.** The rectangles are the largest contigs that cumulatively make up 300 Mbp (~10%) of the assembly: (**a**) for *CSS-BY* and (**b**) for *CSS-SCZ*.

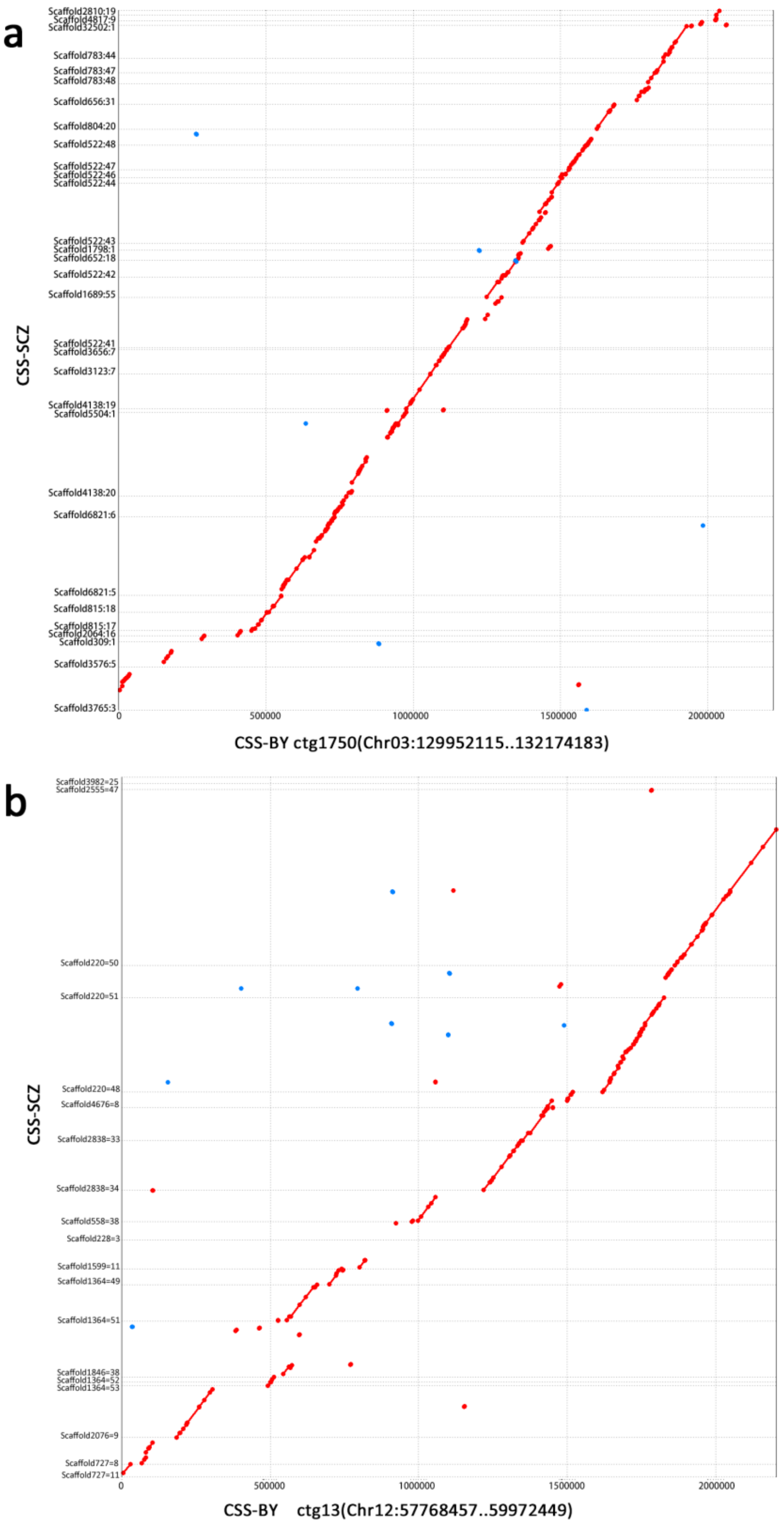


**Supplementary Figure 12. Comparisons of the two assembled tea tree genomes, *CSS-BY* and *CSS-SCZ*, at a contig level. (a)** ctg1750, and **(b)** ctg13. The x-axis shows contigs in *CSS-BY*, and the y-axis indicates the corresponding contigs in *CSS-SCZ*.

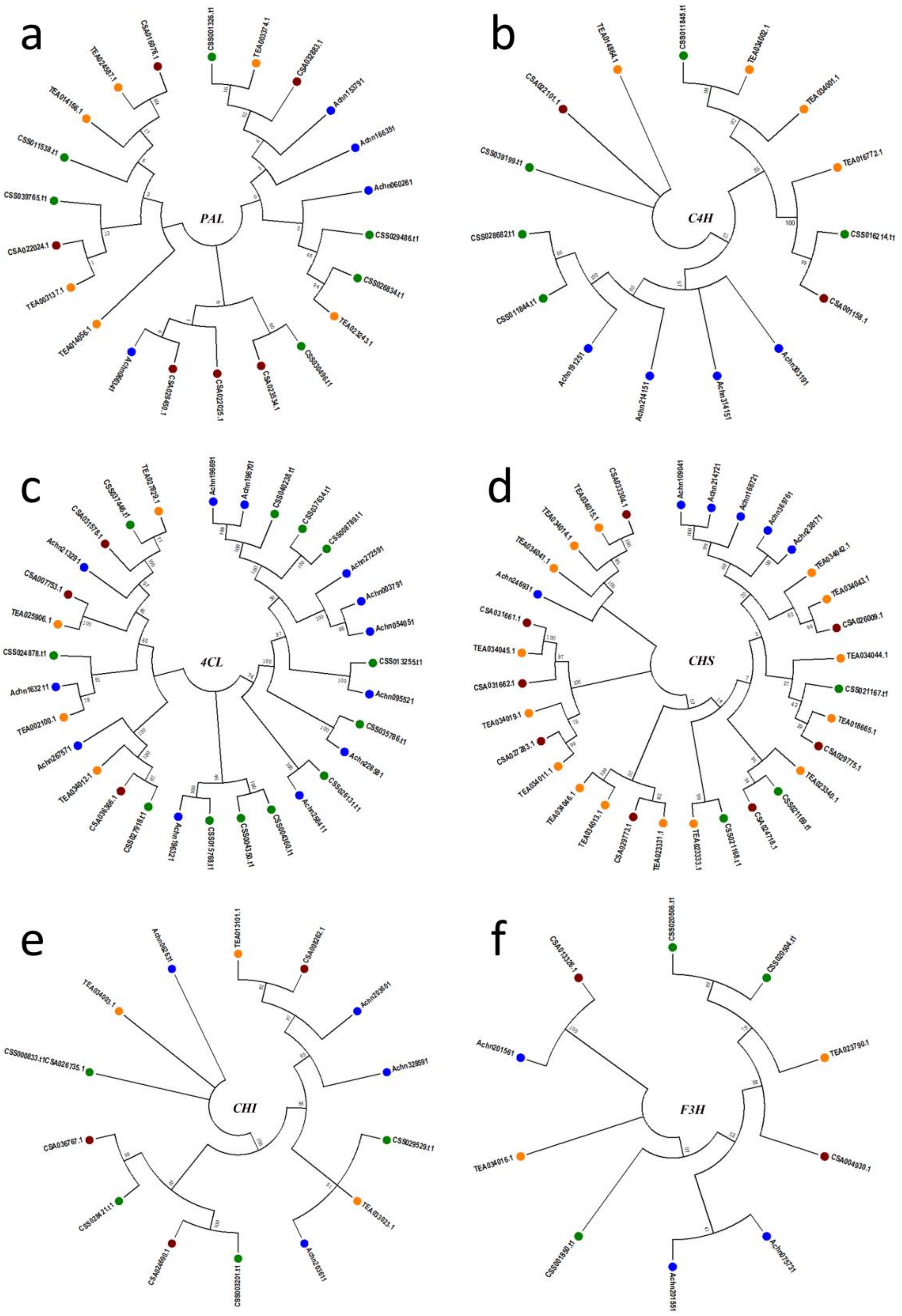
a
PAL
b
C4H
c
4CL
d
CHS
e
CHI
f
F3H

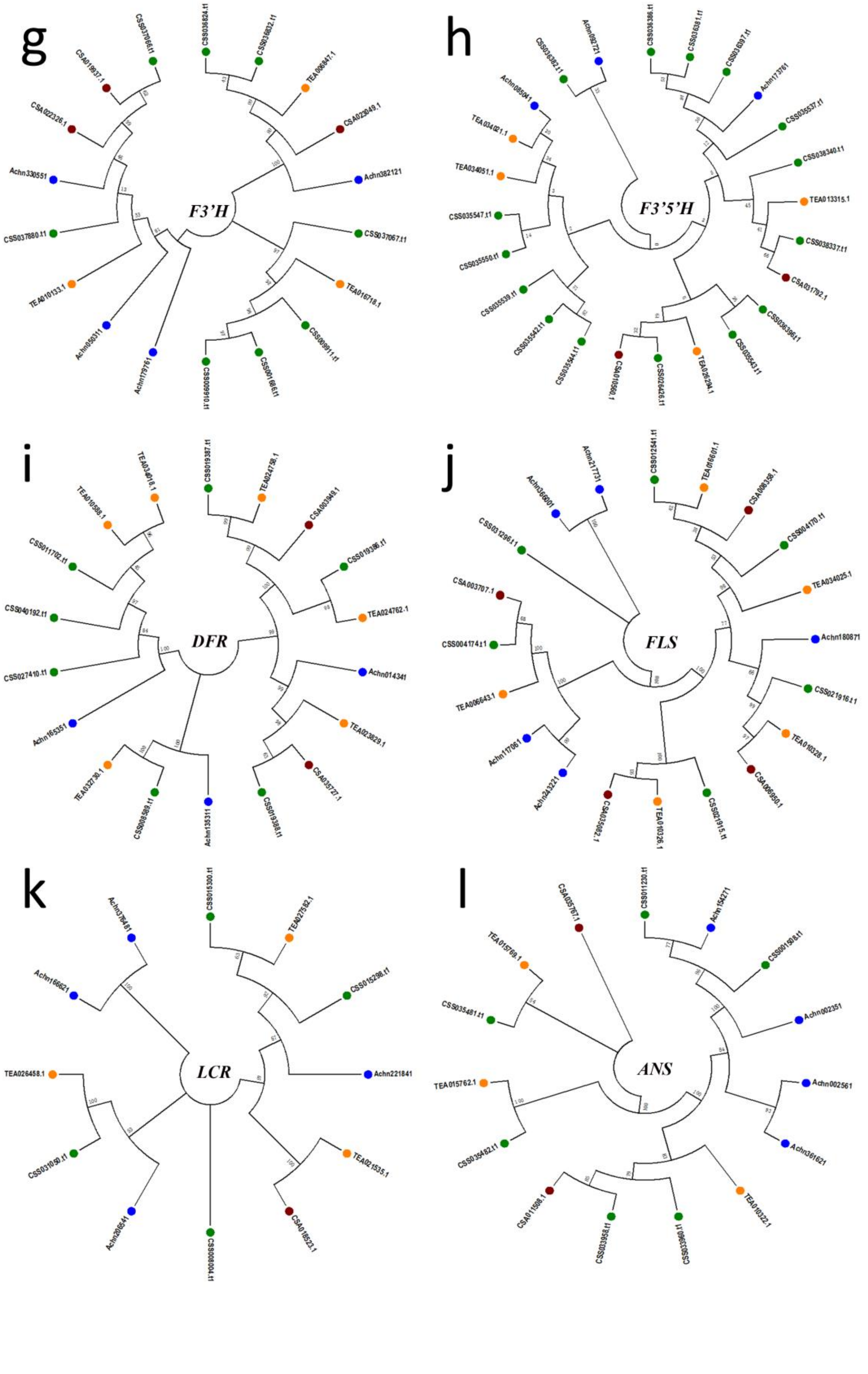
g
F3'H
h
F3'5'H
i
DFR
j
FLS
k
LCR
l
ANS

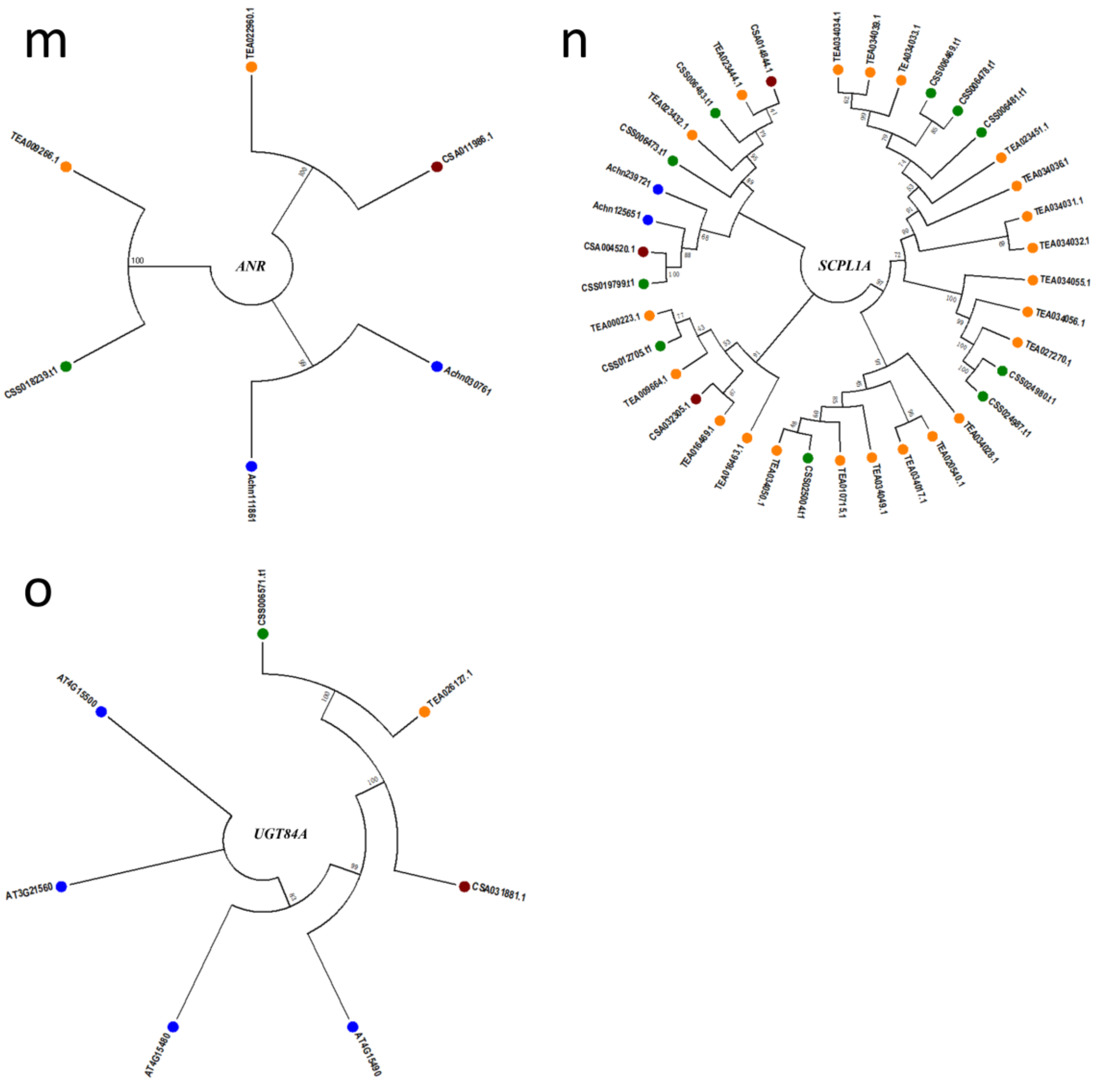


**Supplementary Figure 13. Molecular evolution of 14 major gene families involved in the catechins biosynthesis among three tea tree genome assemblies and kiwifruit. (a)** *PAL*; **(b)** *C4H*; **(c)** *4CL*; **(d)** *CHS*; **(e)** *CHI*; **(f)** *F3H*; **(g)** *F3'H*; **(h)** *F3'5'H*; **(i)** *DFR;* **(j)** *FLS*; **(k)** *LCR*; **(l)** *ANS*; **(m)** *ANR*; **(n)** *SCPL1A* and **(o)** *UGT84A*. Green for *CSS-BY*, orange for *CSS-SCZ*, brown for *CSA-YK10* and blue for kiwifruit.

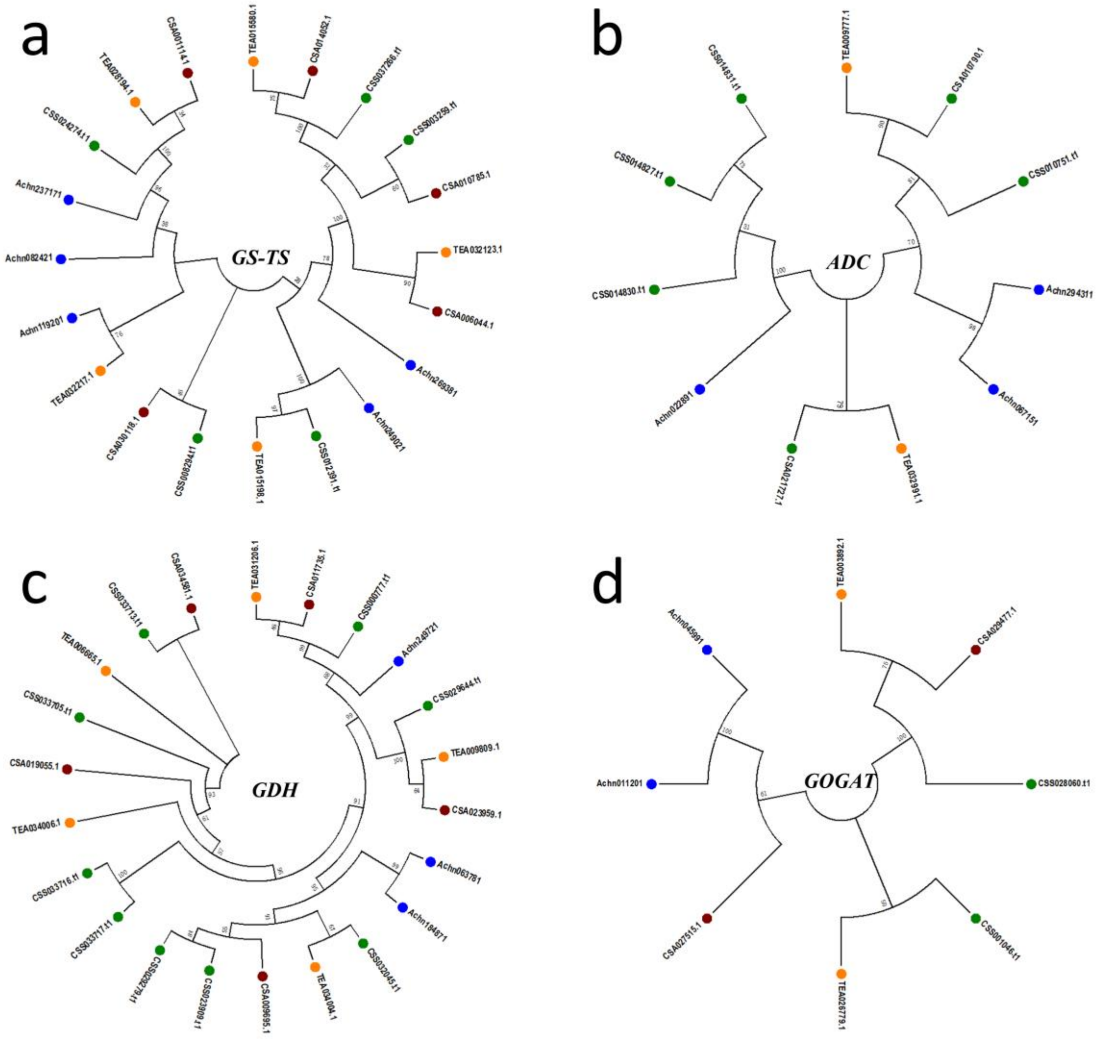


**Supplementary Figure 14. Molecular evolution of the four key gene families involved in the theanine biosynthesis among the three tea tree genome assemblies and kiwifruit. (a)** *GS*/*TS* (putative *TS* genes are shown in pink); **(b)** *ADC*; **(c)** *4GDH*; **(d)** *GOGAT*. Green for *CSS-BY*, orange for *CSS-SCZ*, brown for *CSA-YK10* and blue for kiwifruit.

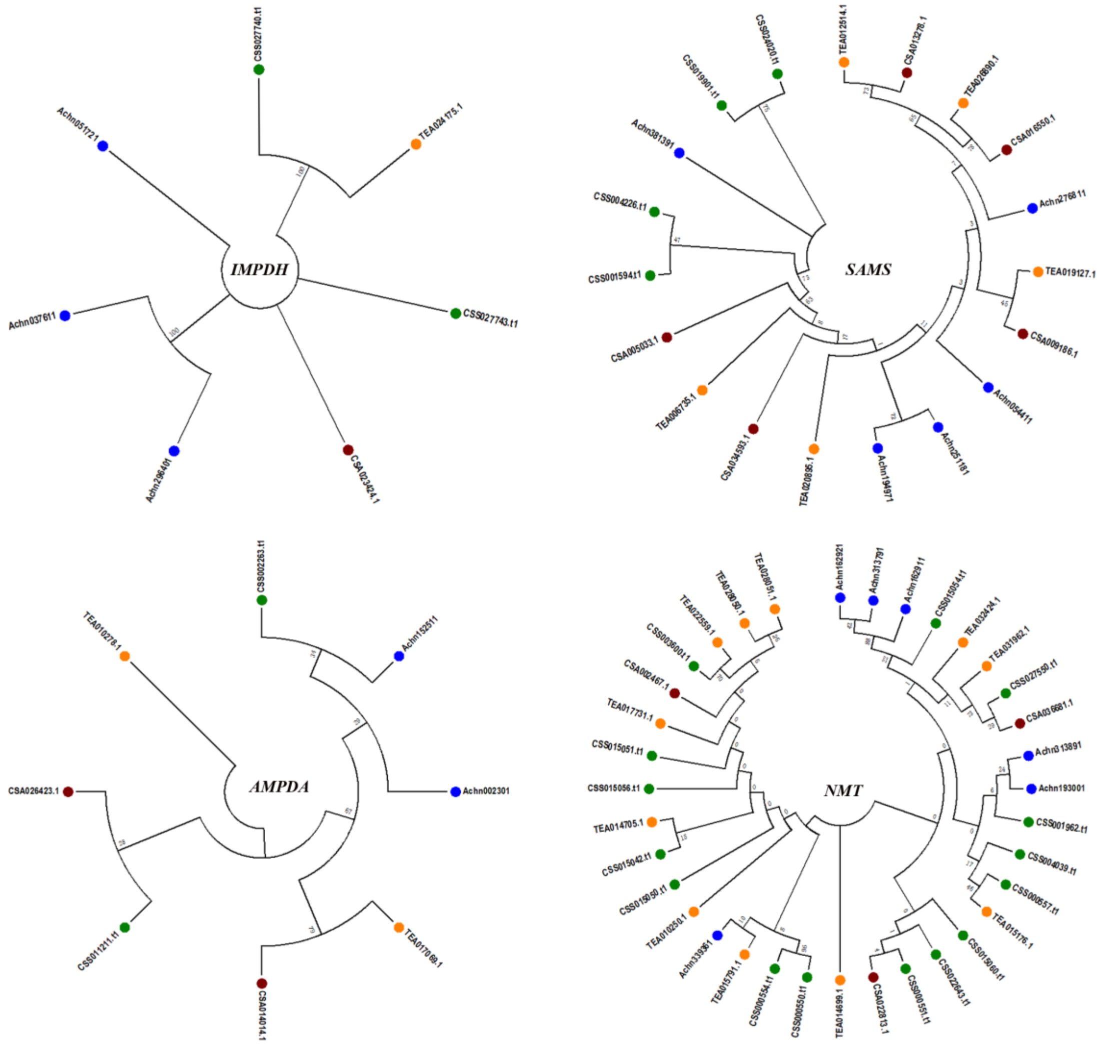


**Supplementary Figure 15. Molecular evolution of the 14 key gene families involved in the caffeine biosynthesis among the three tea tree genome assemblies and kiwifruit. (a)** *IMPDH*; **(b)** *SAMS*; **(c)** *AMPDA* **and (d)** *NMT*. Green for *CSS-BY*, orange for *CSS-SCZ*, brown for *CSA-YK10* and blue for kiwifruit.

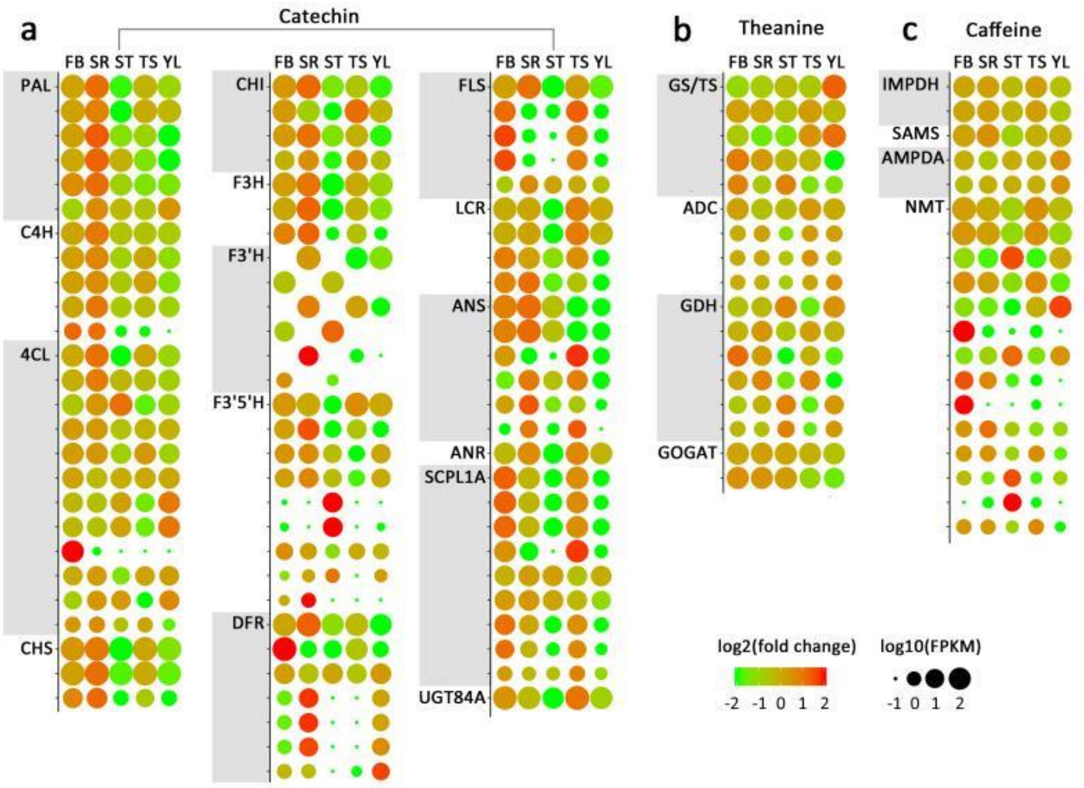


**Supplementary Figure 16.** Expression profiles in FPKM of key functional genes for five tissue related to three metabolic pathways in the tea tree. The candidate genes involved in **(a)** catechin, **(b)** theanine, and **(c)** caffeine biosynthesis pathways. Dot size is plotted as log10 FPKM (fragments per kilobase per million reads mapped), and color for log2 fold-change within columns.

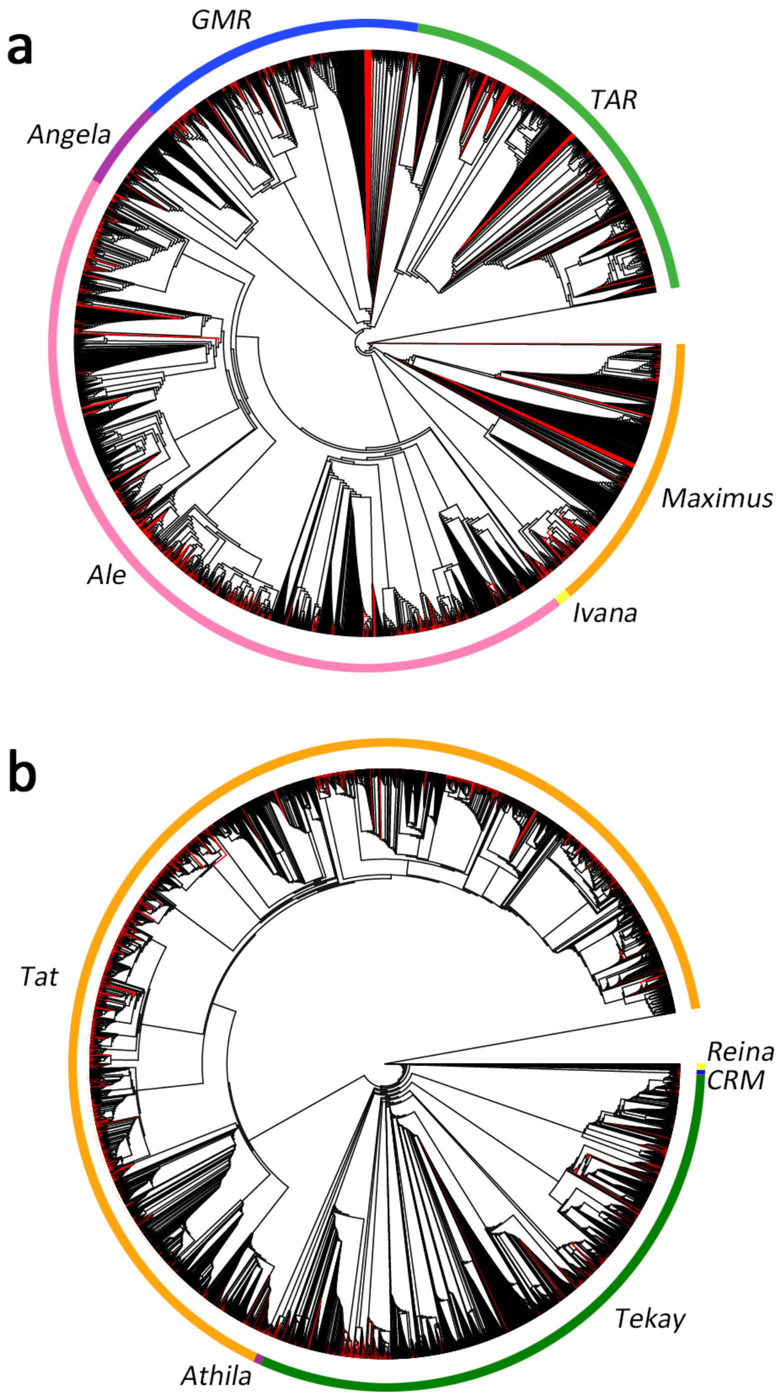


**Supplementary Figure 17. Phylogenetic analyses of LTR retrotransposons of the *C. sinensis* var. *sinensis* and *C. sinensis* var. *assamica* genomes.** The neighbor-joining and unrooted phylogenetic trees were constructed on the basis of 6,036 Ty1-*copia* **(a)** and 17,751 Ty3-*gypsy* **(b)** aligned sequences corresponding to the RT domains without a premature termination codon. The black branch for *CSS-BY* and red branch for *CSA-YK10*.

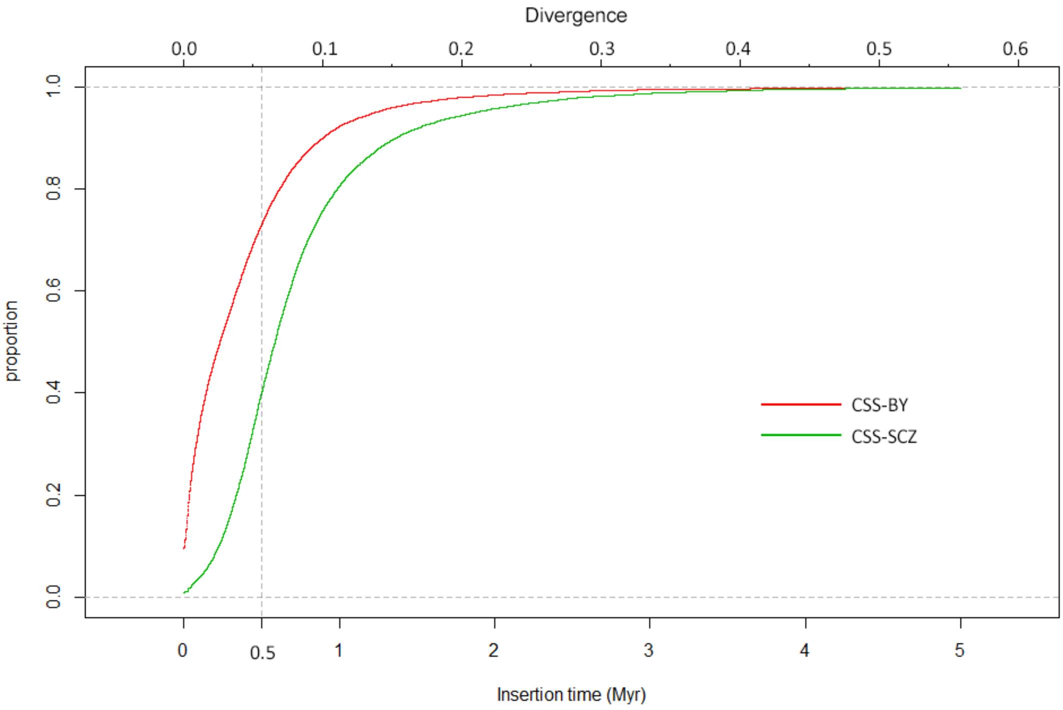


**Supplementary Figure 18. The accumulation curve of LTR retrotransposon insertion times in the *CSS-BY* genome**. The insertion times for LTR retrotransposons were calculated by the formula of $T = K/2r$. T: insertion time; r: synonymous mutations / site / Myr; K: the divergence between the two LTRs. A substitution rate of $5.62\times10^{-9}$ per site per year was used to calculate the insertion times. *CSS-BY* is shown in red, and *CSS- SCZ* is indicated in green. The distributional difference is significant ($p < 2.2e^{-16}$; Wilcoxon rank-sum test) between *CSS-BY* and *CSS- SCZ*.

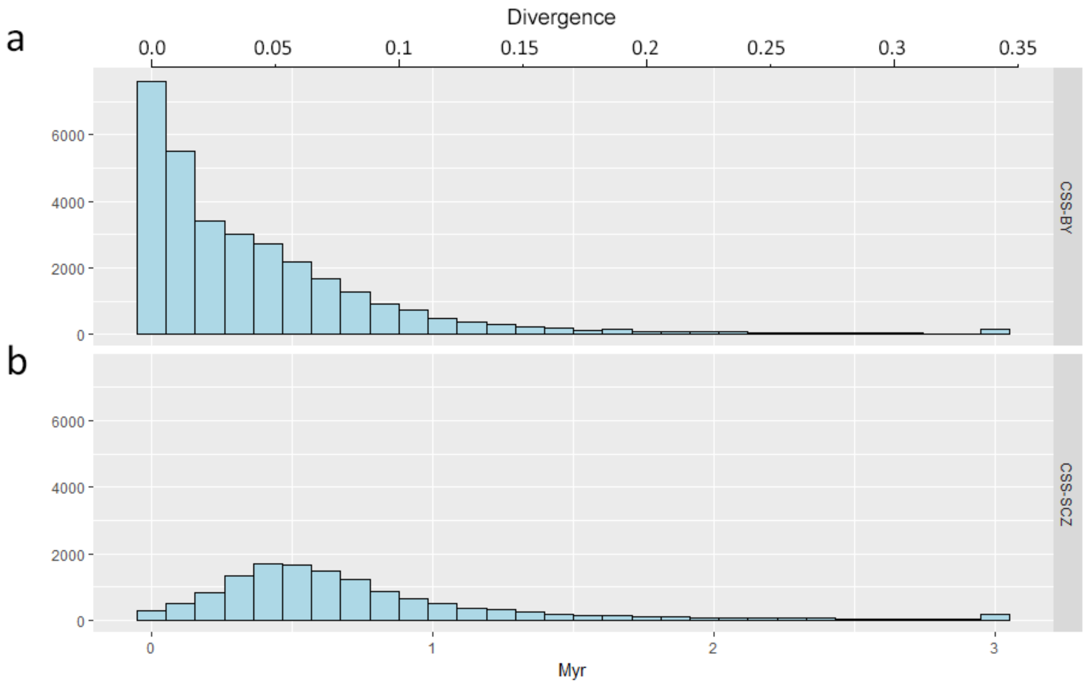


**Supplementary Figure 19. Summary of insertion times of the two tea tree genome assemblies of *C. sinensis* var. *sinensis*. (a)** *CSS-BY*; **(b)** *CSS-SCZ*. The distributional difference is significant ($p < 2.2e^{-16}$; Wilcoxon rank-sum test) between *CSS-BY* and *CSS- SCZ.*

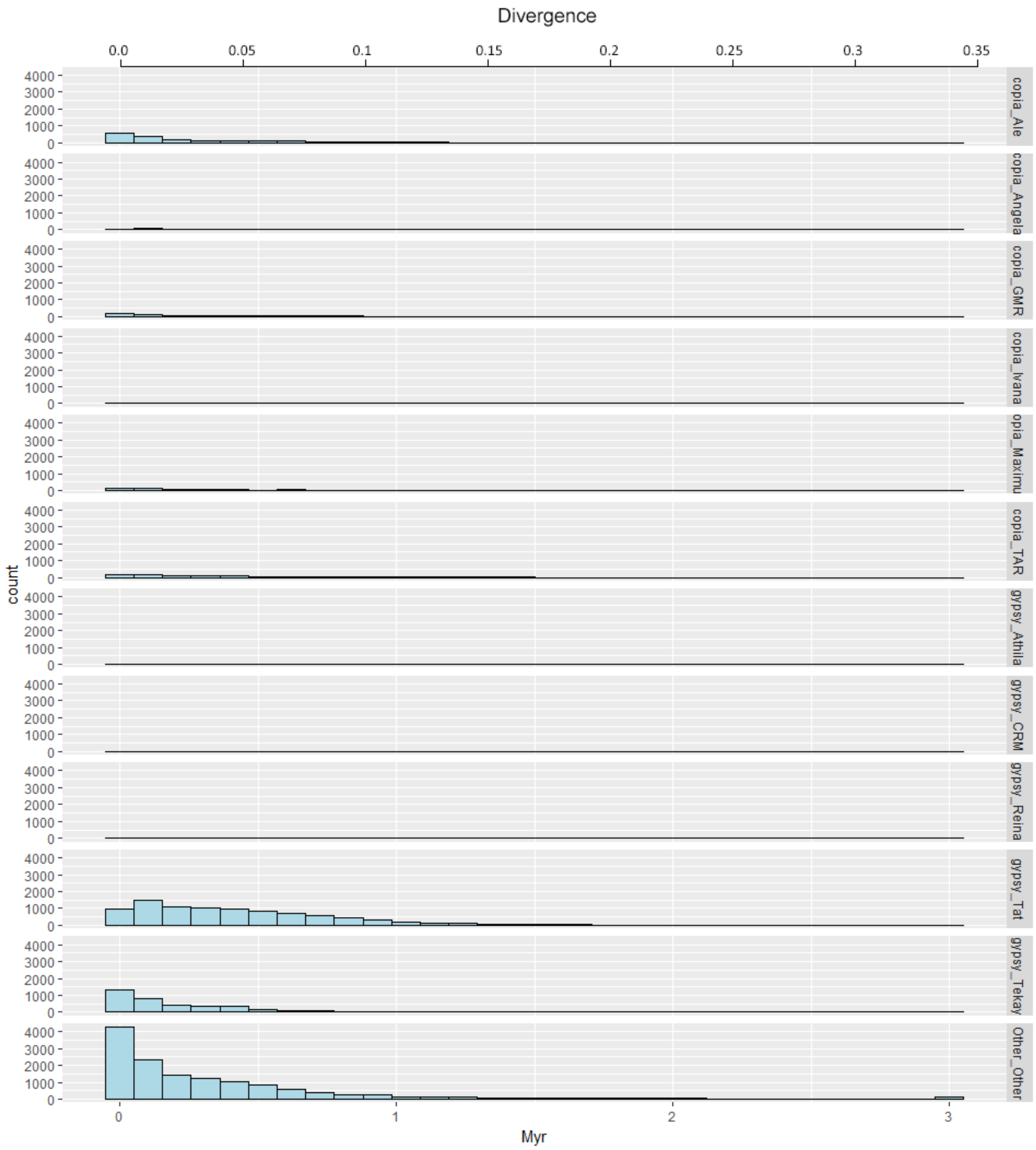


**Supplementary Figure 20. Copy number and insertion times of the six lineages of Ty1-*copia*, five lineages of Ty3-*gypsy* and other types of LTR retrotransposon super-families in the *CSS-BY* genome.**

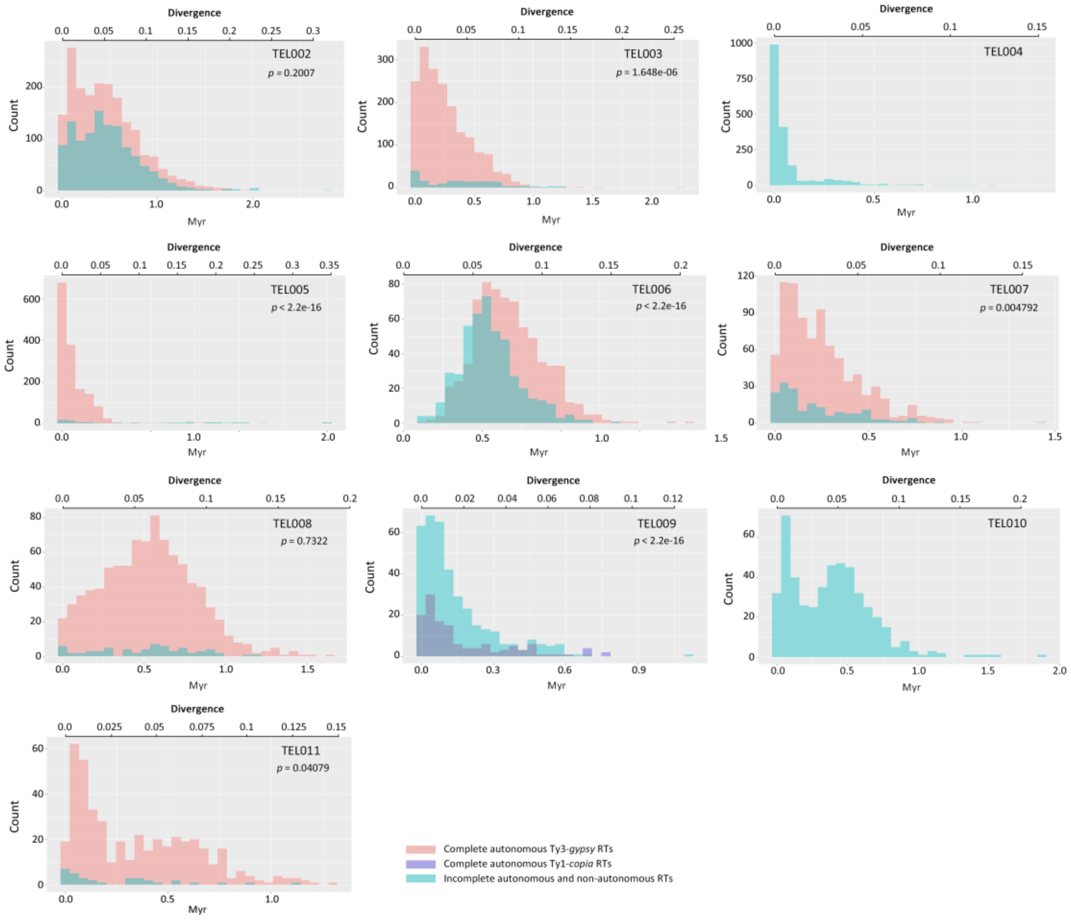


**Supplementary Figure 21. Insertion times of the top 2 to 11 LTR retrotransposon families in the *CSS-BY* genome**. Autonomous Ty1-*copia* retroelements were represented in purple, complete autonomous Ty3-*gypsy* retroelements were represented in red, complete autonomous Ty3-*gypsy* retroelements were represented in purple, while incomplete autonomous and non-autonomous retroelements were represented in blue. The distributional differences were evaluated by Wilcoxon rank-sum test.

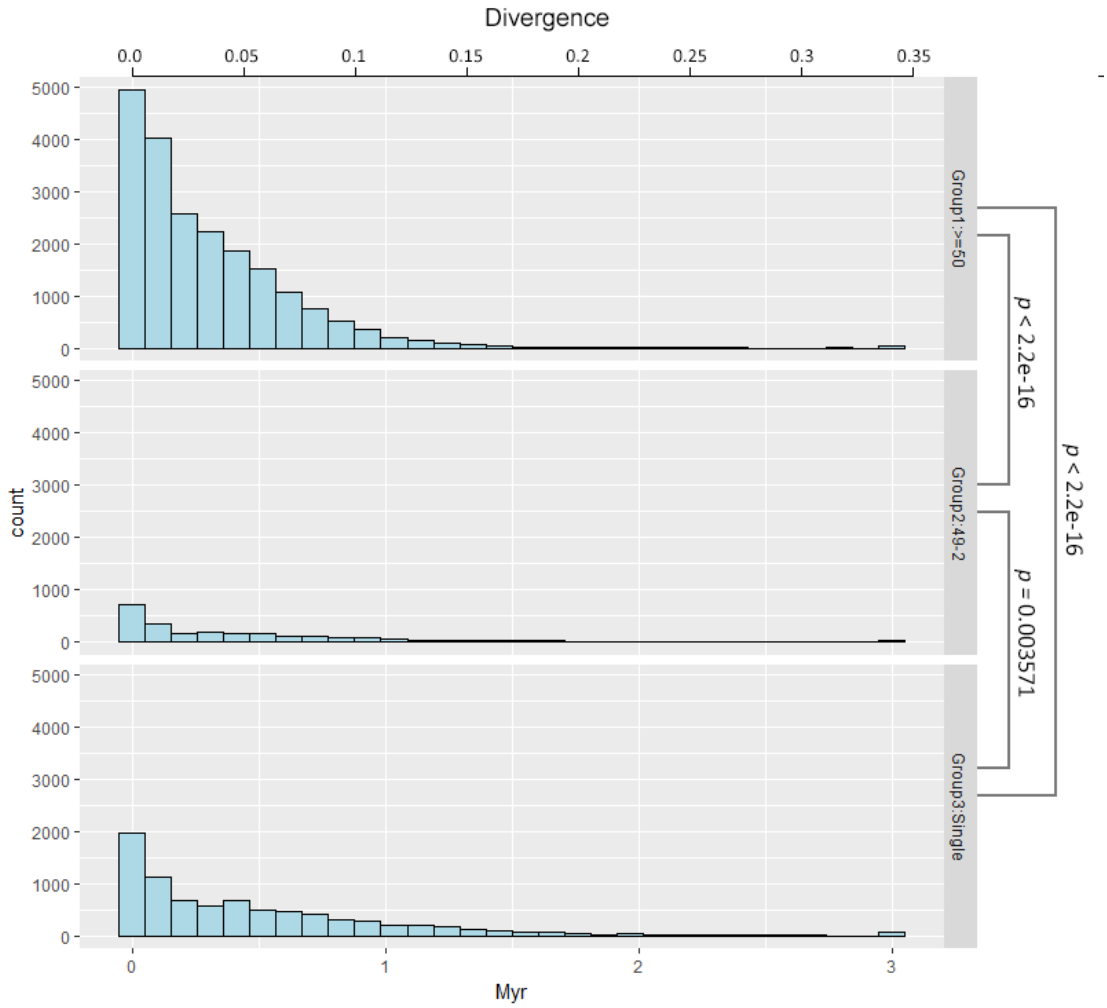


**Supplementary Figure 22. Summary of insertion times of the three types of LTR retrotransposon families in the *C. sinensis* var. *sinensis* cv. *Biyun* genome**. (**a)** high-copy LTR retrotransposon families with >=50 full-length member; (**b)** median-copy LTR retrotransposon families with 2-49 full-length member; (**c)** single-copy LTR retrotransposon families. The distributional differences were evaluated by Wilcoxon rank-sum test.

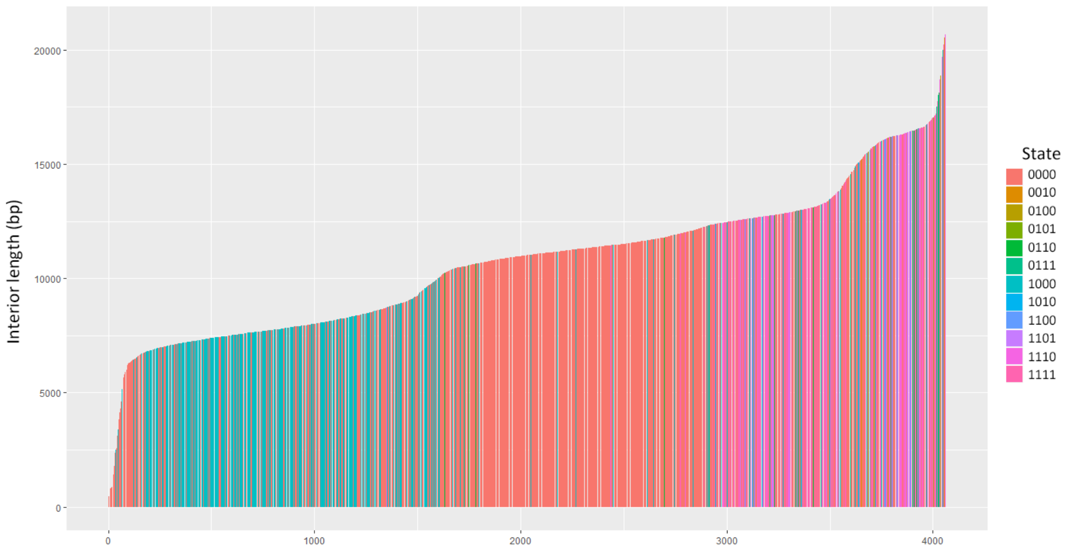


**Supplementary Figure 23. Length distribution of interior regions of the largest LTR retrotransposon family *TEL001*.** The four digits in the 'Description' represent the *gag* and *pol* (PR, RT and IN) genes in LTR retrotransposon, respectively. The number '1' represents the presence of a Pfam annotation, and '0' indicates the absence. For example, '1110' means that the LTR retrotransposon contain a *gag*, PR and RT without IN.

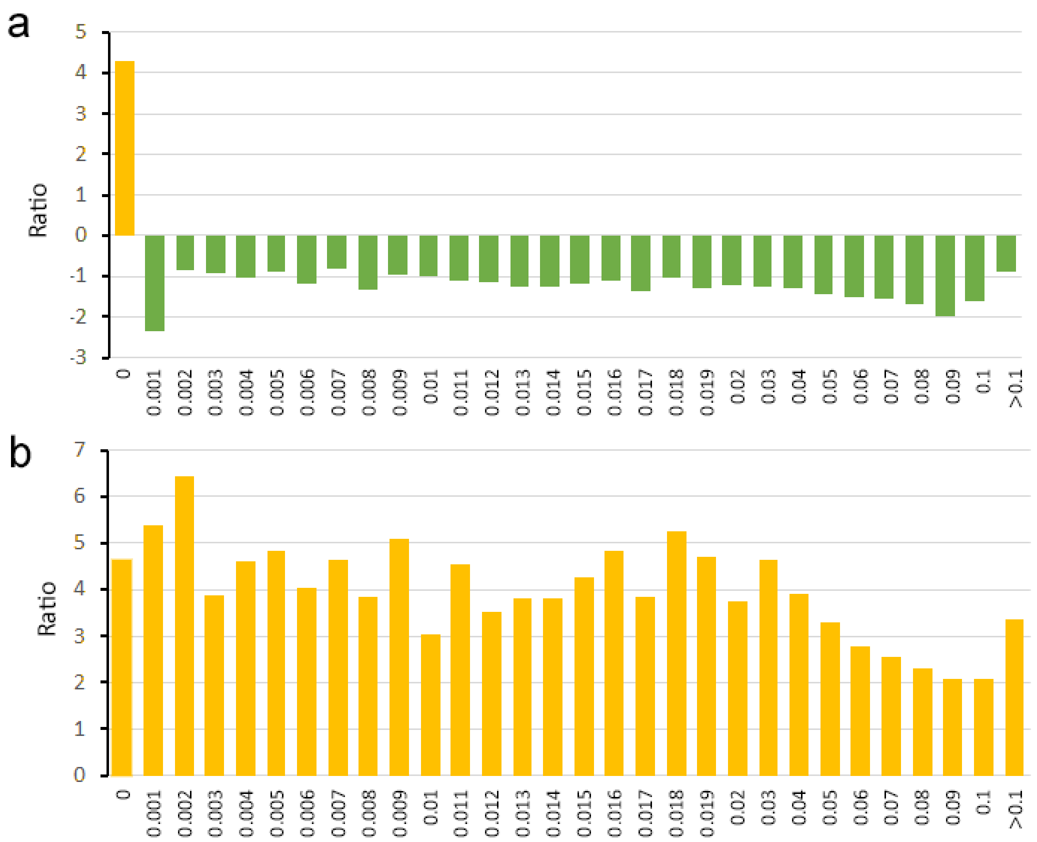


**Supplementary Figure 24. Copy number variation and sequence divergence of Ty1-*copia*, Ty3-*gypsy* and non-autonomous LTR retrotransposons in the tea tree genome.** The X axis represents sequence divergence between the two LTRs, while the Y axis indicates the ratio of copy number of **(a)** non-autonomous / Ty3-*gypsy* retroelements and **(b)** non-autonomous / Ty1-*copia* retroelements. Blue stands for Ty1-*copia*, green stands for Ty3-*gypsy* and yellow stands for non-autonomous LTR retrotransposons, respectively.

### References

Altschul, S.F., Madden, T.L., Schäffer, A.A., Zhang, J., Zhang, Z., Miller, W., and Lipman, D.J. (1997). Gapped BLAST and PSI-BLAST: a new generation of protein database search programs. Nucleic acids research 25:3389-3402.

Anders, S., Pyl, P.T., and Huber, W. (2015). HTSeq--a Python framework to work with high-throughput sequencing data. Bioinformatics 31:166-169.

Ashburner, M., Ball, C.A., Blake, J.A., Botstein, D., Butler, H., Cherry, J.M., Davis, A.P., Dolinski, K., Dwight, S.S., and Eppig, J.T. (2000). Gene Ontology: tool for the unification of biology. Nature genetics 25:25.

Baucom, R.S., Estill, J.C., Leebens-Mack, J., and Bennetzen, J.L. (2009). Natural selection on gene function drives the evolution of LTR retrotransposon families in the rice genome. Genome research 19:243-254.

Boeckmann, B., Bairoch, A., Apweiler, R., Blatter, M.-C., Estreicher, A., Gasteiger, E., Martin, M.J., Michoud, K., O'donovan, C., and Phan, I. (2003). The SWISS-PROT protein knowledgebase and its supplement TrEMBL in 2003. Nucleic acids research 31:365-370.

Burton, J.N., Adey, A., Patwardhan, R.P., Qiu, R., Kitzman, J.O., and Shendure, J. (2013). Chromosome-scale scaffolding of de novo genome assemblies based on chromatin interactions. Nature biotechnology 31:1119.

Cantarel, B.L., Korf, I., Robb, S.M., Parra, G., Ross, E., Moore, B., Holt, C., Alvarado, A.S., and Yandell, M. (2008). MAKER: an easy-to-use annotation pipeline designed for emerging model organism genomes. Genome research 18:188-196.

Chen, C., Xia, R., Chen, H., and He, Y. (2018). TBtools, a Toolkit for Biologists integrating various HTS-data handling tools with a user-friendly interface. BioRxiv:289660.

Chen, N. (2004). Using RepeatMasker to identify repetitive elements in genomic sequences. Current protocols in bioinformatics 5:4.10. 11-14.10. 14.

Delcher, A.L., Salzberg, S.L., and Phillippy, A.M. (2003). Using MUMmer to identify similar regions in large sequence sets. Current protocols in bioinformatics:10.13. 11-10.13. 18.

Dudchenko, O., Batra, S.S., Omer, A.D., Nyquist, S.K., and Hoeger, M. (2017). De novo assembly of the Aedes aegypti genome using Hi-C yields chromosome-length scaffolds. 356:92-95.

Durand, N.C., Robinson, J.T., Shamim, M.S., Machol, I., Mesirov, J.P., Lander, E.S., and Aiden, E.L. (2016). Juicebox provides a visualization system for Hi-C contact maps with unlimited zoom. Cell systems 3:99-101.

English, A.C., Richards, S., Han, Y., Wang, M., Vee, V., Qu, J., Qin, X., Muzny, D.M., Reid, J.G., and Worley, K.C. (2012). Mind the gap: upgrading genomes with Pacific Biosciences RS long-read sequencing technology. PloS one 7:e47768.

Felsenstein, J. (1993). PHYLIP (phylogeny inference package), version 3.5 c: Joseph Felsenstein.

Finn, R.D., Bateman, A., Clements, J., Coggill, P., Eberhardt, R.Y., Eddy, S.R., Heger, A., Hetherington, K., Holm, L., and Mistry, J. (2013). Pfam: the protein families database. Nucleic acids research 42:D222-D230.

Finn, R.D., Tate, J., Mistry, J., Coggill, P.C., Sammut, S.J., Hotz, H.R., Ceric, G., Forslund, K., Eddy, S.R., Sonnhammer, E.L., et al. (2008). The Pfam protein families database. Nucleic Acids Res 36:D281-288.

Grabherr, M.G., Haas, B.J., Yassour, M., Levin, J.Z., Thompson, D.A., Amit, I., Adiconis, X., Fan, L., Raychowdhury, R., and Zeng, Q. (2011). Trinity: reconstructing a full-length transcriptome without a genome from RNA-Seq data. Nature biotechnology 29:644.

Griffiths-Jones, S., Moxon, S., Marshall, M., Khanna, A., Eddy, S.R., and Bateman, A. (2005). Rfam: annotating non-coding RNAs in complete genomes. Nucleic Acids Res 33:D121-124.

Huang, S., Ding, J., Deng, D., Tang, W., Sun, H., Liu, D., Zhang, L., Niu, X., Zhang, X., and Meng, M. (2013). Draft genome of the kiwifruit Actinidia chinensis. Nature communications 4:2640.

Johnson, M., Zaretskaya, I., Raytselis, Y., Merezhuk, Y., McGinnis, S., and Madden, T.L. (2008). NCBI BLAST: a better web interface. Nucleic acids research 36:W5-W9.

Jones, P., Binns, D., Chang, H.-Y., Fraser, M., Li, W., McAnulla, C., McWilliam, H., Maslen, J., Mitchell, A., and Nuka, G. (2014). InterProScan 5: genome-scale protein function classification. Bioinformatics 30:1236-1240.

Kanehisa, M., and Goto, S. (2000). KEGG: kyoto encyclopedia of genes and genomes. Nucleic acids research 28:27-30.

Lagesen, K., Hallin, P., Rodland, E.A., Staerfeldt, H.H., Rognes, T., and Ussery, D.W. (2007). RNAmmer: consistent and rapid annotation of ribosomal RNA genes. Nucleic Acids Res 35:3100-3108.

Larkin, M.A., Blackshields, G., Brown, N.P., Chenna, R., McGettigan, P.A., McWilliam, H., Valentin, F., Wallace, I.M., Wilm, A., Lopez, R., et al. (2007). Clustal W and Clustal X version 2.0. Bioinformatics 23:2947-2948.

Li, H., and Durbin, R. (2009). Fast and accurate short read alignment with Burrows–Wheeler transform. bioinformatics 25:1754-1760.

Li, L., Stoeckert, C.J., and Roos, D.S. (2003). OrthoMCL: identification of ortholog groups for eukaryotic genomes. Genome research 13:2178-2189.

Llorens, C., Futami, R., Covelli, L., Dominguez-Escriba, L., Viu, J.M., Tamarit, D., Aguilar-Rodriguez, J., Vicente-Ripolles, M., Fuster, G., Bernet, G.P., et al. (2011). The Gypsy Database (GyDB) of mobile genetic elements: release 2.0. Nucleic Acids Res 39:D70-74.

Lowe, T.M., and Eddy, S.R. (1997). tRNAscan-SE: a program for improved detection of transfer RNA genes in genomic sequence. Nucleic Acids Res 25:955-964.

Lowe, T.M., and Eddy, S.R. (1999). A computational screen for methylation guide snoRNAs in yeast. Science 283:1168-1171.

McCarthy, E.M., and McDonald, J.F. (2003). LTR_STRUC: a novel search and identification program for LTR retrotransposons. Bioinformatics 19:362-367.

Möller, S., Croning, M.D., and Apweiler, R. (2001). Evaluation of methods for the prediction of membrane spanning regions. Bioinformatics 17:646-653.

Nawrocki, E.P., Kolbe, D.L., and Eddy, S.R. (2009). Infernal 1.0: inference of RNA alignments. Bioinformatics 25:1335-1337.

Parra, G., Bradnam, K., and Korf, I. (2007). CEGMA: a pipeline to accurately annotate core genes in eukaryotic genomes. Bioinformatics 23:1061-1067.

Pertea, M., Kim, D., Pertea, G.M., Leek, J.T., and Salzberg, S.L. (2016). Transcript-level expression analysis of RNA-seq experiments with HISAT, StringTie and Ballgown. Nature protocols 11:1650.

Roach, M.J., Schmidt, S.A., and Borneman, A.R. (2018). Purge Haplotigs: allelic contig reassignment for third-gen diploid genome assemblies. BMC bioinformatics 19:460.

Robinson, J.T., Turner, D., Durand, N.C., Thorvaldsdottir, H., Mesirov, J.P., and Aiden, E.L. (2018). Juicebox. js provides a cloud-based visualization system for Hi-C data. Cell systems 6:256-258. e251.

SanMiguel, P., Gaut, B.S., Tikhonov, A., Nakajima, Y., and Bennetzen, J.L. (1998). The paleontology of intergene retrotransposons of maize. Nat Genet 20:43-45.

Servant, N., Varoquaux, N., Lajoie, B.R., Viara, E., Chen, C.J., Vert, J.P., Heard, E., Dekker, J., and Barillot, E. (2015). HiC-Pro: an optimized and flexible pipeline for Hi-C data processing. Genome Biol. 16:259.

Shi, T., Huang, H., and Barker, M.S. (2010). Ancient genome duplications during the evolution of kiwifruit (Actinidia) and related Ericales. Annals of Botany 106:497-504.

Simão, F.A., Waterhouse, R.M., Ioannidis, P., Kriventseva, E.V., and Zdobnov, E.M. (2015). BUSCO: assessing genome assembly and annotation completeness with single-copy orthologs. Bioinformatics 31:3210-3212.

Smit, A., Hubley, R., and Green, P. (2016). RepeatMasker Open-4.0. 2015. Google Scholar.

Tamura, K., Dudley, J., Nei, M., and Kumar, S. (2007). MEGA4: Molecular Evolutionary Genetics Analysis (MEGA) software version 4.0. Mol Biol Evol 24:1596-1599.

Vitte, C., Panaud, O., and Quesneville, H. (2007). LTR retrotransposons in rice (*Oryza sativa*, L.): recent burst amplifications followed by rapid DNA loss. BMC Genomics 8:218.

Walker, B.J., Abeel, T., Shea, T., Priest, M., Abouelliel, A., Sakthikumar, S., Cuomo, C.A., Zeng, Q., Wortman, J., and Young, S.K. (2014). Pilon: an integrated tool for comprehensive microbial variant detection and genome assembly improvement. PloS one 9:e112963.

Wang, Y., Tang, H., DeBarry, J.D., Tan, X., Li, J., Wang, X., Lee, T.-h., Jin, H., Marler, B., and Guo, H. (2012). MCScanX: a toolkit for detection and evolutionary analysis of gene synteny and collinearity. Nucleic acids research 40:e49-e49.

Wei, C., Yang, H., Wang, S., Zhao, J., Liu, C., Gao, L., Xia, E., Lu, Y., Tai, Y., and She, G. (2018). Draft genome sequence of Camellia sinensis var. sinensis provides insights into the evolution of the tea genome and tea quality. Proceedings of the National Academy of Sciences:201719622.

Wicker, T., Sabot, F., Hua-Van, A., Bennetzen, J.L., Capy, P., Chalhoub, B., Flavell, A., Leroy, P., Morgante, M., and Panaud, O. (2007). A unified classification system for eukaryotic transposable elements. Nature Reviews Genetics 8:973.

Xia, E.-H., Zhang, H.-B., Sheng, J., Li, K., Zhang, Q.-J., Kim, C., Zhang, Y., Liu, Y., Zhu, T., and Li, W. (2017). The tea tree genome provides insights into tea flavor and independent evolution of caffeine biosynthesis. Molecular plant 10:866-877.

Yang, Z. (2007). PAML 4: phylogenetic analysis by maximum likelihood. Mol Biol Evol 24:1586-1591.

Yao, X., Tang, P., Li, Z., Li, D., Liu, Y., and Huang, H. (2015). The first complete chloroplast genome sequences in Actinidiaceae: genome structure and comparative analysis. PloS one 10:e0129347.
